# 3D-printed polymeric scaffolds with optimized architecture to repair a sheep metatarsal critical-size bone defect

**DOI:** 10.1101/2022.12.14.520447

**Authors:** Charlotte Garot, Sarah Schoffit, Cécile Monfoulet, Paul Machillot, Claire Deroy, Samantha Roques, Julie Vial, Julien Vollaire, Martine Renard, Hasan Ghanem, Hanane El-Hafci, Adeline Decambron, Véronique Josserand, Laurence Bordenave, Georges Bettega, Marlène Durand, Mathieu Manassero, Véronique Viateau, Delphine Logeart-Avramoglou, Catherine Picart

**Affiliations:** Université de Grenoble Alpes, CEA, INSERM U1292 Biosanté, CNRS EMR 5000 Biomimetism and Regenerative Medicine (BRM), 17 avenue des Martyrs, F-38054 Grenoble, France; Ecole Nationale Vétérinaire d’Alfort, Université Paris-Est, F-94704 Maisons-Alfort, France; Université Paris Cité, CNRS, INSERM, ENVA, B3OA, F-75010 Paris, France; Univ. Bordeaux, INSERM, Institut Bergonié, CIC 1401, F-33000 Bordeaux, France; CHU de Bordeaux, CIC-IT, INSERM, Institut Bergonié, CIC 1401, F-33000 Bordeaux, France; INSERM U1209, Institute of Advanced Biosciences, F-38000 Grenoble, France; Université Grenoble Alpes, Institute of Advanced Biosciences, F-38000 Grenoble, France; Service de chirurgie maxillo-faciale, Centre Hospitalier Annecy Genevois, 1 avenue de l’hôpital, F–74370 Epagny Metz-Tessy, France; Institut Universitaire de France

**Author notes:** Co-second authors (Sarah Schoffit, Cécile Monfoulet, Paul Machillot, Claire Deroy, Samantha Roques, Julie Vial). Co-third authors (Julien Vollaire, Martine Renard, Hasan Ghanem, Hanane El-Hafci). Co-corresponding authors (Marlène Durand), (Véronique Viateau), (Delphine Logeart-Avramoglou), (Catherine Picart).

**Keywords:** tissue engineering, 3D printing, medical devices, surface coatings, bone, large animal model

## Abstract

The reconstruction of critical-size bone defects in long bones remains a challenge for clinicians. We developed a new bioactive medical device for long bone repair by combining a 3D-printed architectured cylindrical scaffold made of clinical-grade polylactic acid (PLA) with a polyelectrolyte film coating delivering the osteogenic bone morphogenetic protein 2 (BMP-2). This film-coated scaffold was used to repair a sheep metatarsal 25-mm long critical-size bone defect. *In vitro* and *in vivo* biocompatibility of the film-coated PLA material were proved according to ISO standards. Scaffold geometry was found to influence BMP-2 incorporation. Bone regeneration was followed using X-ray scans, µCT scans, and histology. We showed that scaffold internal geometry, notably pore shape, influenced bone regeneration, which was homogenous longitudinally. Scaffolds with cubic pores of ∼870 µm and a low BMP-2 dose of ∼120 µg/cm^3^ induced the best bone regeneration without any adverse effects. The visual score given by clinicians during animal follow-up was found to be an easy way to predict bone regeneration. This work opens perspectives for a clinical application in personalized bone regeneration.

## 1. Introduction

Critical-size bone defects, which result from high energy traumas, non-unions, tumor resection, infection, etc. are unable to heal by themselves [1]. Thus, their reconstruction remains a challenge for clinicians and has a high economical and societal cost [2]. Currently, autologous bone graft is the gold standard solution to treat such defects, but it is associated with some drawbacks such as limited availability, increase in the number of surgeries required, significant postoperative donor-site morbidity, and inconsistency of repair in large bone defects exceeding typically about 5 cm^3^ [3,4]. To tackle these drawbacks, synthetic scaffolds, made of a large variety of materials, including ceramics, metals, and polymers, have been developed and used as bone graft substitutes [5–7]. For efficient bone regeneration, these scaffolds should be biocompatible, preferably biodegradable and bioresorbable with appropriate kinetics, osteoconductive, osteoinductive, and possess sufficient mechanical properties [5]. To date, scaffolds are widely used to replace or consolidate bones in maxillofacial surgery, dentistry, spine and orthopedics. However, the use of scaffolds alone is not sufficient to manage challenging clinical situations. The induction of bone regeneration can be achieved by optimizing the properties of the scaffolds, or by including additional functionalities, such as active surface coatings [8– 13], delivery of osteoinductive growth factors like bone morphogenetic proteins [14–19], or stem cells [15,20,21]. Such strategies are currently being developed in view of clinical applications [22].

In terms of regulation, stem cell-based products are categorized as “advanced medicinal therapeutic products”, which requires a strong control of their source, culture conditions prior to implantation, and knowledge of their fate once implanted *in vivo* [23]. Growth factor-or drug-based products, i.e. without stem cells, need to follow the regulatory path for medicinal products, pharmaceutical products, or combined medical devices, depending on their intended use and whether the scaffold itself is responsible or not of the main action [22,24,25]. Bone morphogenetic proteins 2 and 7 (BMP-2 and BMP-7) are the most widely used osteoinductive growth factors for bone regeneration and, combined with a collagen sponge or paste, have been clinically approved for the treatment of open fractures of long bones, non-unions, and spinal fusion [26,27]. Unfortunately, some controversies arose because supra-physiological doses (several milligrams) of these proteins were used for their application and have resulted in adverse effects in humans. In fact, the use of collagen sponge, which has a poor affinity for BMPs, results in massive protein leakage into the soft tissue followed by systemic diffusion leading to the occurrence of inflammation, osteolysis, bone cyst, and ectopic bone formation [28]. More recently, new forms of BMP proteins, such as the BV-265 chimera, have been developed and new materials have been engineered to optimize and localize the delivery of BMPs [29]. Two major strategies are used to optimize their delivery in space and time. First, delivering BMPs via a hydrogel or a paste. In this case, BMPs are usually pre-mixed with the hydrogel or paste, which is then either directly injected into the bone defect, or used to fill a synthetic scaffold with mechanical properties suitable for bone regeneration and the combination is inserted into the bone defect. For instance, BMP-7 was loaded into a collagen carrier that was inserted in the inner duct of a composite synthetic scaffold made of polycaprolactone (PCL) and tricalcium phosphates (β-TCP) [15,30]. Second, BMPs may be incorporated into a surface coating [32,33], which is itself deposited at the surface of a synthetic scaffold. This approach enables to control both the mechanical and physico-chemical properties of the scaffold and the bioactivity of the BMP protein delivered via the surface coating. Being localized at the scaffold surface, the delivery of the protein is spatially controlled. Surface coatings made by layer-by-layer assembly of polyelectrolytes and loaded with BMPs have already been used to control BMP dose at an implant surface [33,34]. We previously evidenced that a film based on hyaluronic acid (HA) and poly(L-lysine) (PLL) can deliver tunable doses of BMP-2 to repair a critical-size bone defect in a rat femur [35]. Promisingly, such films can be dried and sterilized before implantation, while remaining osteoinductive [36].

Recent developments in material engineering have open new possibilities in custom material design and fabrication. Due to their versatility, additive manufacturing (AM) techniques have recently emerged and can be applied to the fabrication of 3D scaffolds made of metals like titanium, ceramics like β-TCP and hydroxyapatite, or polymers [7,37]. More specifically, fused deposition modeling (FDM) is the most widely used AM technique to fabricate synthetic polymeric scaffolds of any size and shape [37].

In order to be clinically translated, synthetic grafts must be fabricated using good manufacturing practice-grade raw components and processes. Biocompatibility assays need to be performed *in vitro* according to regulatory standards, notably NF EN ISO 10993, to ensure that the new products meet the safety requirements. Following these *in vitro* normative tests, *in vivo* biocompatibility and biodegradability assays can be performed in small animal models, again following the NF EN ISO 10993 guidelines.

Scaffold geometry should be designed to support and guide bone formation within large bone defects. As a result, scaffolds should be porous and provide sufficiently interconnected pore space for fluid flow and cell invasion, particularly before the completion of angiogenesis, in order to promote bone growth inside the pores [38]. However, the ideal internal architecture features of scaffolds, including the porosity, pore size and shape, pore connectivity, surface area, surface convexities/concavities, etc. have not been evidenced yet, since they depend on several parameters, especially the implantation site. Several studies have focused on the optimization of scaffold geometry but the vast majority of them were conducted *in vitro* and results depended on the experimental conditions [39,40]. The difficulty also arises from the interdependency between parameters. For instance, some studies focused on pore size, but tuning this parameter also modifies porosity and scaffold stiffness. Thus, it was difficult to conclude on the independent effect of the studied parameter. The few *in vivo* studies focused mainly on pore size and suggested conflicting results [41–47]. These discrepancies may be explained by the differences in animal models, implantation sites, volume of the defect (non-critical versus critical-size), scaffold materials, fabrication methods, and pore shapes. The vast majority of these studies were performed in small animal models (mice, rats, and rabbits). To our knowledge, only one study was conducted in a large animal model, i.e. sheep [48]. Globally, it is generally agreed that pore size ranging between 300 and 800 µm is preferred [39]. To our knowledge, the influence of pore shape on bone regeneration has barely been studied *in vivo* [49–51]. These studies showed that orthogonal pores led to more bone formation compared to hexagonal pores and gyroid pores induced more bone formation than orthogonal pores [49–51]. The type of cubic unit cell may also influence bone regeneration [51].

Preclinical experiments *in vivo* should preferably be performed in large animal models that best mimic human bone physiology. Minipig, dog, and sheep are the most widely used large animal models for bone regeneration [15,37,52]. Minipigs are often used for mandibular bone defects [53], while sheep are rather used for orthopedic bone defects, mimicking the localization of tibial fractures. Furthermore, bone metabolism of sheep is similar to that in humans [1,54–56]. In a recent study, we showed that a 3D-printed polymeric scaffold coated with the (PLL/HA) biomimetic film containing a controlled dose of BMP-2 was effective to repair a very large (12 cm^3^) critical-size bone defect in the mandible of minipigs [53]. The local environment in experimental mandibular bone defects provides a large amount of well-vascularized cortico-cancellous bone with extensive muscle coverage, which creates conditions conducive to BMP-2-induced bone regeneration. The regeneration of long bone segmental defects still need to be proved using such bioactive film-coated synthetic scaffolds.

In the present study, we aimed at developing a bioactive synthetic bone graft substitute to repair a critical-size metatarsal bone defect in sheep, which could be subsequently translated to the clinic. To this end, we designed and fabricated 3D-printed scaffolds made of polylactic acid (PLA) and we coated them with an osteoinductive (PLL/HA) film containing a controlled dose of BMP-2. First, the biocompatibility and biodegradability of the film-coated PLA were assessed *in vitro* and *in vivo* in a small animal model according to the regulatory standards for implantable biomaterials. Then, 3D scaffolds with different scaffold internal architectures were designed, fabricated, and assessed for their efficiency to repair a 25 mm-long sheep metatarsal bone defect. Together, our results suggest that biocompatible and biodegradable 3D-printed PLA scaffolds with cubic pores of ∼870 µm coated with a biomimetic film and loaded with a controlled BMP-2 dose can repair a sheep metatarsal critical-size bone defect. This opens perspectives for clinical applications of such scaffolds that can be personalized to each patient.

## 2. Materials and methods

### 2.1 Materials and chemicals

Polyethyleneimine (PEI, Sigma, Saint-Quentin-Fallavier, France), sodium hyaluronate (HA, Lifecore, Chaska, USA), and poly(L-lysine) (PLL, Sigma) were dissolved respectively to 5 mg/mL, 1 mg/mL, and 0.5 mg/mL in a N-(2-hydroxyethyl)piperazine-N′-ethanesulfonic acid (HEPES)/NaCl buffer (20 mM HEPES at pH 7.4, 0.15 M NaCl). HEPES and NaCl were purchased from Sigma. 1-ethyl-3-(3-dimethylaminopropyl)carbodiimide (EDC, Sigma) and N-hydroxysulfosuccinimide (Sulfo-NHS, Chemrio, Ningbo, China) were dissolved to 30 or 50 mg/mL for EDC (respectively abbreviated EDC30 and EDC50) and 11 mg/mL for Sulfo-NHS in 0.15M NaCl at pH 5.5. BMP-2 was obtained from InductOs kit (Medtronic, USA). PLL labeled with fluorescein 5-isothiocyanate (PLL^FITC^) and 5(6)-Carboxytetramethylrhodamine N-succinimidyl ester (Rhod) were purchased from Sigma.

Clinical-grade PLA (cgPLA, Lactoprene 100M Monofilament, Poly-Med, Inc, Anderson, USA) was used for BMP-2 loading experiments and all *in vivo* experiments and PLA filament of regular-grade (rgPLA, Verbatim, PLA filament 1.75mm, Düsseldorfer, Germany) was used for other experiments.

### 2.2. Design and 3D printing of PLA scaffolds

The external shape of the scaffolds was designed using OnShape (http://www.onshape.com). Then, the internal architecture was designed on Ultimaker Cura 4.5 (Ultimaker B.V., Utrecht, Netherlands) by choosing an infill pattern corresponding to the pore shape (either Cubic for cubic opened pores, Zigzag for cubic semi-closed pores or Gyroid for gyroid pores) and an infill density or infill line distance corresponding to the targeted pore size. This architecture was then modified by directly modifying the code piloting the 3D printer, named G-code. This G-code was modified manually and using a homemade Python code. Once designed, the scaffolds were 3D-printed using an Ender-3 3D printer (Creality3D®, Shenzhen, China). Different types of 2D PLA discs and 3D scaffolds were designed and 3D-printed depending on the purpose of each specific *in vitro* or *in vivo* experiment (**Figure 1**). For all types of scaffolds, PLA filaments deposited layer-by-layer were ∼400 µm large and ∼200 µm in height. The 3D-architectured scaffolds had always an external geometry consisting in a cylinder with a differentiated inner ring. Their unit cell was either cubic, gyroid, or a combination of cubic and gyroid. The 2D discs were not architectured.

**FIGURE 1.**
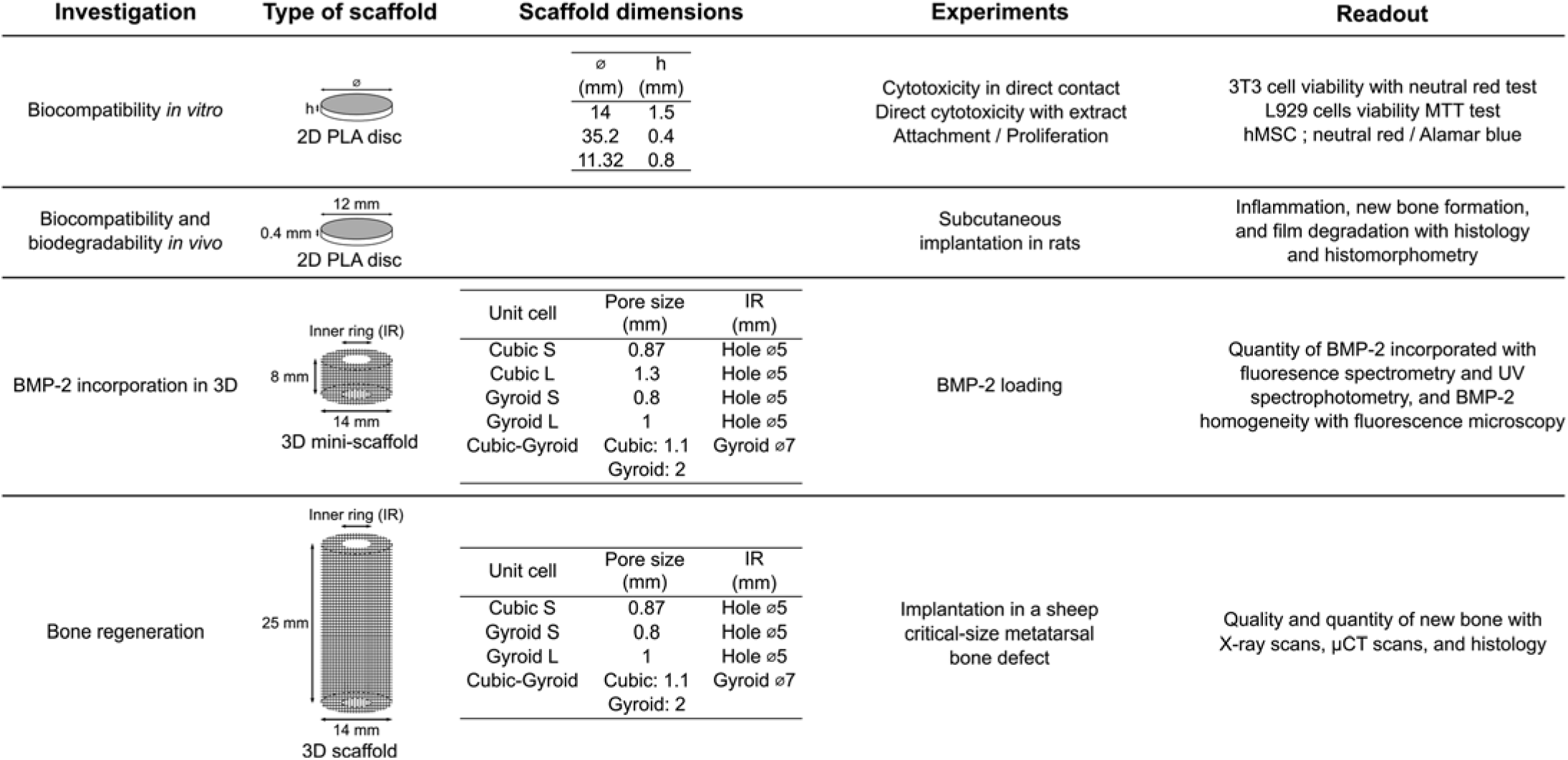
Summary of the experimental design. The different investigations led are specified, along with the types of scaffolds used for each investigation, their dimensions, the experiments performed, and the readouts of the experiments.

### 2.3. Designs of experiments (DOE)

DOE were performed using a dedicated software (Design-Expert® 12, Stat-Ease, Inc., Minneapolis, USA) to select the scaffold geometries to implant in a sheep critical-size metatarsal bone defect (**Table SI 1**). The factors (parameters to be optimized) of the DOE were the infill density and infill pattern of scaffolds. The readouts were the mechanical properties (compressive modulus and compressive strength), the porosity (expressed in %), and the effective pore size. To select the geometries, porosity and mechanical properties were maximized and pore size was targeted to be large enough (> 800 µm) to allow adequate vascular and bone ingrowth. This was a factorial design that was randomized, the design type was Plackett Burman, and response modeling was reduced to main effects.

### 2.4. Film coating of PLA scaffolds and BMP-2 loading

Prior to film coating, PLA scaffolds were always pre-wetted in ultrapure water for 24 h. Polyelectrolyte multilayer films were deposited at the surface of 2D PLA discs fabricated for *in vitro* biocompatibility assays, except the 35.2 mm diameter discs, with a liquid-handling robot (EVO100, Tecan, Lyon, France) as described by Machillot et al. [57]. The 35.2 mm diameter PLA discs, PLA discs for *in vivo* experiments on rats, and 3D PLA mini-scaffolds and scaffolds were coated using a dip-coating robot (Dipping Robot DR 3, Riegler & Kirstein GmbH, Potsdam, Germany) as described previously [35,58]. Briefly, a first layer of PEI was deposited. Then, 24 alternating layer pairs of PLL and HA were deposited to form a (PLL/HA)_24_ film. Films were subsequently crosslinked using EDC and Sulfo-NHS as previously described [35,53]. EDC50 was used for *in vitro* experiments except BMP-2 incorporation assays and EDC30 was used for *in vivo* experiments and BMP-2 incorporation tests. Scaffolds were finally rinsed with HEPES/NaCl buffer. BMP-2 was post-loaded in the films. For biocompatibility *in vitro* and biocompatibility and biodegradability *in vivo* experiments, two different concentrations were used: 30 µg/mL (low dose, BMP-2 LD) and 60 µg/mL (high dose, BMP-2 HD). For BMP-2 incorporation in 3D experiments, the BMP-2 concentration varied between 10 and 50 µg/mL, named hereafter BMP10, BMP30, and BMP50. For bone regeneration experiments, BMP-2 was post-loaded in the film at a targeted dose of ∼500 µg/total volume of defect, based on our previous study led on minipigs [53]. We took into account the fact that scaffolds had different surfaces and volumes and adapted the BMP-2 concentration in the loading solution to achieve the targeted BMP-2 dose. Thus, BMP-2 concentrations in the loading solutions were 43.2 µg/mL for Cubic S, 27.9 µg/mL for Gyroid L, 34.8 µg/mL for Cubic-Gyroid, and 38.2 µg/mL for Gyroid S. BMP-2 was incubated for 2 h at 37°C. PLA scaffolds were then rinsed before being dried under a biological safety cabinet. For biocompatibility *in vitro* experiments, prepared samples were stored in multiwell plates sealed with Parafilm® (Sigma-Aldrich, Saint-Quentin-Fallavier, France) and stored at 4°C. Before beginning the assays, the plates were γ-sterilized at 25 kGy for 92 h (Gamma Cell 3000 Elan, MDS Nordion Canada). For all other experiments, scaffolds were UV-sterilized after preparation.

### 2.5. In vitro biocompatibility assays

Four biocompatibility assays were performed on the 2D PLA discs following ISO 10993 part 5 guidelines [59]: i) direct cytotoxicity with contact: 3T3 cells (Balb 3T3, clone A31, ATCC) were cultured in a medium with serum (DMEM and 10% SV, ATCC) and put in direct contact with the biomaterials during 24 h (*n* = 4 for all conditions). A qualitative evaluation was done by microscopy and a quantitative evaluation was done by measuring cell viability with neutral red that targeted lysosomal activity; ii) direct cytotoxicity with extract: L929 cells (NCTL L929, ATCC) were cultured without serum and set in contact for 24 h with extracts and extract dilutions of biomaterials (*n* = 4 for all conditions). Then, a qualitative evaluation was performed by microscopy and a quantitative evaluation was done by measuring cell viability with MTT test (Sigma) that targeted mitochondrial activity. For these two cytotoxicity assays, thermanox (Nunc) was used as negative control and latex as positive control; iii) attachment: 50,000 human mesenchymal stem cells (hMSC, Promocell) were cultured without serum (only 2% SV) and set in contact with the biomaterials in 48-well microplates (*n* = 3 for all conditions). A quantitative evaluation was done after 15 h by a dosage of lysosomal activity (p-nitrophenyl-n-acetyl-betaD-glucosamide, Sigma-Aldrich); and iv) proliferation: 5,000 hMSC were cultured in mesenchymal stem cell growth medium (Promocell) and set in contact with the biomaterials in 48-well microplates (*n* = 3 for all conditions). A quantitative evaluation was then performed with Alamar blue test at different time points: 1, 4, 7, 11, and 14 days. For these two latest assays, the control was cells cultured directly on plastic. For each of these assays, four experimental conditions were chosen: bare PLA, film-coated PLA without BMP-2, and film-coated PLA with BMP-2 loaded at two different BMP-2 concentrations: BMP-2 LD and BMP-2 HD. The dimensions of the discs used for the different assays are specified in **Figure 1**.

### 2.6. In vivo subcutaneous implantation of 2D PLA discs in rats for assessment of in vivo biocompatibility and biodegradability of polyelectrolyte films

Following ISO 10993-2, thirty-six 8-week old Wistar rats (Charles River, France) weighing ∼250 g were included in the study, which was approved by the animal ethics committee (*APAFIS#33421-20211101211184565 v3*). They were acclimated for 10 days before surgery. Before anesthesia, perioperative analgesia was implemented using buprenorphine at 0.1 mg/kg by intraperitoneal injection. Animals were then anesthetized via an induction cage (5% isoflurane and 2 L/min oxygen), and the anesthesia was maintained using a mask (2.5% isoflurane and 2 L/min oxygen). An ophthalmic gel (Ocrygel, TVM) was used during anesthesia and put in place before animal clipping. The animals were placed in prone position before clipping and disinfecting their back skin using chlorinetetracycline. Hypothermia was avoided by heating the induction cages and the surgical plan at 37°C using a heating mat. Following ISO 10993-6 recommendations, the PLA discs of 12 mm in diameter and 0.4 mm in height (**Figure 1**) were implanted subcutaneously. For that, 4 incisions of 1 cm were made in the back region and near the flanks using an 11 mm blade. A subcutaneous detachment was made with Metzembaum scissors. A prepared PLA disc was placed in the created pocket. The created pockets were not interconnected. The wound was then closed using intradermal suture with a 4-0 monofilament (polyglecaprone 25). After surgery, analgesia was maintained when necessary by subcutaneous injection of buprenorphine (0.02 mg/kg). During recovery from anesthesia and until their awakening, the rats were placed in heated cages. A qualified technician clinically evaluated the animals every day, except during the weekend, for the first 7 days after surgery. After the first 7 days, animals were clinically evaluated once a week. Their general condition, feeding, watering, cleaning, morbidity, and mortality was observed and taken care of. Animals were weighed once a week until euthanasia. The study was composed of 4 experimental groups: bare PLA, film-coated PLA without BMP-2, and film-coated PLA with BMP-2 loaded at two different BMP-2 concentrations: BMP-2 LD and BMP-2 HD. All experimental groups were studied at three time points: day 7, 28, and 48 (D7, D28, D48), with *n* = 3 rats for each experimental condition at each time point. The animals were euthanized by pentobarbital intraperitoneal injection followed by an injection of Exagon® (1 mL/kg) combined with lidocaine (10 mg/mL). The materials were removed with surrounding tissues and samples from spleen, kidney, brain, heart, and liver. All these specimens were preserved and stored in 4% paraformaldehyde at 4°C until analysis.

### 2.7. Histology and histomorphometry of 2D PLA discs

Specimens were fixed in 4% paraformaldehyde formalin (Antigenfix, Microm Microtech, France) with stirring for 24 h. Dehydration of specimens was conducted in ascending alcohol baths (Absolute ethanol ≥ 99.8%, VWR Chemicals), then in toluene (Toluene N/A ≥ 99%, VWR Chemicals), followed by paraffin (Paraplast X-TRA®, LEICA Biosystems) impregnation using a LEICA TP1020. They were then embedded in paraffin using a LEICA EG1150H embedding platform. Sections ≤ 6 µm were cut using a microtome (LEICA RM2255) and stained with Hematoxylin Erythrosine Saffron staining (HES). They were then analyzed with a microscope (Nikon NiU). Histomorphometry was conducted using a dedicated software (NIS Elements D). To quantify the film still present at the implant surface after explantation and sample processing, the percentage of the film remaining at the PLA discs surface was calculated based on the measurements of the disc perimeters and of the film length along the disc surface.

### 2.8. Study of BMP-2 loading in 3D

To assess the effect of pore size and shape on BMP-2 loading inside different 3D architectures, cylinders of 8 mm in height and 14 mm in diameter, hereafter called 3D mini-scaffolds, were designed and 3D-printed (**Figure 1**) with cgPLA. Five scaffold architectures were used: Cubic S, Cubic L, Gyroid S, Gyroid L, and Cubic-Gyroid. After coating with the polyelectrolyte film, BMP-2 was loaded at BMP10, BMP30, and BMP50 (*n* = 3 3D mini-scaffolds per geometry and per BMP-2 concentration). The homogeneity of BMP-2 loading inside the film-coated 3D mini-scaffolds was assessed using a BMP-2 loading solution containing 5% of BMP-2^Rhod^ and a fluorescence macroscope (Macrofluo Z16 Apo, Leica Microsystems, Wetzlar, Germany) with a 0.8X objective. BMP-2 loading was quantified using a fluorescence spectrometer (Tecan Spark, Tecan Lyon, France) for BMP10 (excitation at 535 nm, bandwidth 25 nm; emission at 595 nm, bandwidth of 35 nm; number of flashes was set at 30 and the integration time at 40 µs) and a UV-Visible spectrophotometer (Cary 60, Agilent Technologies, Inc., Santa Clara, USA) for BMP30 and BMP50. The amount of loaded BMP-2 was measured by quantifying the initial concentration of BMP-2 in the loading solution and the remaining BMP-2 concentration after loading in the 3D mini-scaffolds. The amount of BMP-2 loaded was calculated as the difference between these two concentrations multiplied by the volume of the BMP-2 loading solution.

### 2.9. Characterization of scaffolds and film coating

Two types of PLA filaments were used in this study: rgPLA, which was used for all *in vitro* experiments except the BMP-2 incorporation tests in 3D, and cgPLA, which was used for all *in vivo* experiments and the BMP-2 incorporation tests in 3D. The differences between the two filaments, notably their crystallinity that influences the degradation rate, were investigated by attenuated total reflectance Fourier transform infrared spectroscopy (ATR-FTIR) and small angles X-ray scattering (SAXS). A Ge crystal, Perkin-Elmer spectroscope, and Spectrum software were used for ATR-FTIR. Pieces of PLA filaments were used for the measurements, and background was subtracted for every measurement. Spectra were collected in the range of 4000 to 600 cm^-1^, at a 2 cm^-1^ resolution and with 16 scans. For the SAXS acquisition, PLA filaments were cut and the pieces of filaments were arranged in order to create 1 cm^2^ squares. Acquisitions were made in θ/2θ reflection mode and cobalt radiation (λ=0.17903 nm) was used.

3D mini-scaffolds and 3D scaffolds were imaged by micro-computed tomography (µCT) using a VivaCT 40 (SCANCO Medical AG, Brüttisellen, Switzerland) to quantify their porosity and surface area. The acquisition parameters were set at 70 kV with an intensity of 114 µA, an isotropic voxel size of 76 µm, and integration time of 100 ms. For 3D scaffolds, 4 geometries were characterized: Cubic S, Gyroid S, Gyroid L, and Cubic-Gyroid. 3D mini-scaffolds were imaged using an Ultra 55 scanning electron microscope (SEM, Zeiss, Oberkochen, Germany). Prior to imaging, 3D mini-scaffolds were metallized with Platinum. Pore sizes were evaluated at 10 kV with a secondary electron (SE2) detector. The mechanical properties of the 3D scaffolds were measured by performing uniaxial compressive tests with a traction machine (MTS Systems Corporation, Eden Prairie, USA). A10 kN load cell at a speed of 1 mm/s was used. Tests were performed until 10% of deformation was reached. Tests were done in triplicate for each scaffold geometry. The compressive strength and compressive modulus (both expressed in MPa) were deduced from the stress-strain curves. The compressive strength was defined as the maximum stress withstood by the scaffolds and the compressive modulus was defined as the slope of the linear (elastic) part of the stress-strain curve.

To image the film deposited onto the 3D scaffolds, the last deposited layer was labeled with FITC (PLL^FITC^) [34,53]. The scaffolds were then stored in 0.15 M NaCl until imaging using a fluorescence macroscope (Macrofluo Z16 Apo, Leica Microsystems, Wetzlar, Germany) with a 0.8X objective.

### 2.10. In vivo sheep metatarsal critical-size bone defect

Twenty-four mature female Pré-Alpes sheep, mean age of 36.2 months (21.2-61.7 months) and weighing 63 kg (46.5-79 kg) were included in this study, which was approved by the animal ethics committee (*APAFIS#20287-2019041715086916 v2*). The animals were obtained from “Les élevages Christian Lebeau” (Gambais, France). Animal housing and care were carried out using procedures previously described [54]. The pre-surgical (notably anesthesia) and surgical procedures were performed as previously described [52]. Briefly, a 25-mm long mid-diaphyseal osteotomy was performed in the left metatarsal bone with full periosteal removal. This defect was stabilized with an osteosynthesis plate (3.5 Dynamic Compression Plate, Synthes) and cortical screws of 3.5 mm in diameter. An implant was then inserted into the defect. Cerclages using two 2/0 polydioxanone sutures (PDS® I, Ethicon) were used around the implant and plate to enhance stability at the replacement site. The wound was then closed. A cylindrical cast including a 5-mm-diameter steel walking bar was placed around the operated hind limb of each sheep. Aftercare was conducted as previously described [52]. The animals were followed up for 4 months. A veterinarian clinically evaluated the animals every day.

Each implant was randomized, implanted, and analyzed in a blind manner using X-ray s. In the preliminary experiment, three scaffold geometries were tested (*n* = 2): Cubic S, Gyroid L, and Cubic-Gyroid, all coated with the biomimetic film and loaded with BMP-2. In the main experiment, two selected conditions were further studied in larger groups to perform statistical analysis. The Cubic S geometry loaded with BMP-2 (*n* = 5), hereafter referred to as Cubic S + BMP-2, was kept and the Gyroid S geometry loaded with BMP-2 (*n* = 7), referred to as Gyroid S + BMP-2, was introduced in order to compare two scaffold geometries with a similar pore size (870 µm and 805 µm for Cubic S and Gyroid S, respectively). Two negative controls were added for each geometry: film-coated scaffolds without BMP-2, referred to as Cubic S w/o BMP-2 and Gyroid S w/o BMP-2 (see **Table 1** for all experimental conditions). Additional groups using autografts (standard care) or defects left empty were not added in the present study, as the authors previously provided evidence of consistent occurrence of bone union using autologous bone grafting and absence of union in an empty defect in the same model in sheep [60]. It was a way to reduce the number of animals used, which is in line with the 3Rs principle on animal research. The two Cubic S + BMP-2 scaffolds implanted in the preliminary experiment were included in the analysis of the main experiment to reach *n* = 7 without the need to use more sheep. Sheep were euthanized after 4 months by overdose of barbiturate. The cast of each sheep was removed and the left metatarsus was excised, radiographed, and fixed in 4% paraformaldehyde under mild shaking for 2 weeks.

**Table 1.** Experimental conditions and total number of sheep studied per experimental group. For the preliminary experiment, *n* = 2 for each condition. For the main experiment, *n* = 2 Cubic S w/o BMP-2 and Gyroid S w/o BMP-2 were added as negative controls (no BMP-2), *n* = 5 Cubic S + BMP-2 were added, and *n* = 7 Gyroid S + BMP-2 were added. Film was crosslinked at EDC30 and loaded with BMP-2. In this table, the first number on the left of the “+” refers to the conditions of the preliminary experiment, while the second number refers to the main experiment.

| Scaffold geometry | Condition |  |
| --- | --- | --- |
|  | Film | Film |
|  | No BMP-2 | + BMP-2 |
| Cubic S | 0 + 2 | 2 + 5 |
| Gyroid S | 0 + 2 | 0 + 7 |
| Gyroid L | - | 2 + 0 |
| Cubic-Gyroid | - | 2 + 0 |

### 2.11. Analysis of bone growth within 3D scaffolds

Qualitative assessment of bone formation was done by X-ray scans each month, and after bone resection at 4 months post-surgery. These radiographs were acquired using an EvolutX FP veterinary radiograph system (Medec Loncin, Belgium) under anesthesia. The acquisition parameters were set to 69 kV and 12.8 mAs. A score, adapted from the one previously developed by the authors, was used to qualify bone formation [53]:

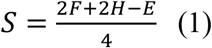

This score took into account three criteria: the percentage of filling of the porous implant (F), the homogeneity of the newly formed bone (H), and the amount of bone outside of the implant, i.e. ‘ectopic’ bone (E). Each criterion was evaluated by five clinicians in a blind manner using a score between 0 and 4. The score was represented as the mean score ± standard deviation (SD) of the scores given by each evaluator independently.

After fixation of the explants, the osteosynthesis plates were removed and the defect sites, along with 1 cm of the surrounding host bone on each edge, were collected and stored in water. Specimens were then imaged with a high-resolution µCT (Skyscan1172, Bruker) with the following settings: 90 kV source voltage, 279 mA source current, 17.7 µm pixel size, 0.3° rotation step, 420 ms exposure time, a frame averaging of 8, and aluminum-copper filters. The scanned images were reconstructed as a stack of slices of each sample using NRecon software (V1.7.4.6, Bruker). A 3D reconstruction of samples was made using CT Vox software (CT Vox v.3.3.1, Bruker). The reconstructed images were then imported into Dragonfly software (ORS Inc., Canada) for quantitative analyses with binarization threshold for bone determined by Otsu’s method. For quantification of the total newly formed bone between edges, data were treated with a volume of interest corresponding to a cylinder centered in the middle of the defect with a length equal to the (25 mm) defect one. The VOI was also divided into three equal parts corresponding to the proximal, central, and distal areas.

### 2.12. Undecalcified histology

Specimens were dehydrated and embedded in methyl methacrylate resin as described in Viateau et al. [52]. The embedded specimens were cut along the metatarsal axis using a circular saw (200-300 µm, Leitz 1600, Leica Biosystems) and a central section (closest to the mid-sagittal plane) and a peripheral section were selected for histological analysis, ground down to 100 µm thick, polished, and stained with Stevenel blue and Van Gieson picrofushin.

### 2.13. Statistical analysis

Origin 2020 (OriginLab Corporation) and Excel (Microsoft Office) were used for all graphical and statistical analyses. Data were expressed as mean ± SD. Non-parametric data were presented by median and interquartile range. Differences between groups were assessed by analysis of variance and Bonferroni post-hoc analysis or Student’s t-test for parametric data, and by Kruskal-Wallis ANOVA with Dunn’s test, Dunnett’s test, and Mann-Whitney U test for non-parametric data. Two-way ANOVA was used to assess differences when two independent variables were used. G* power 3.1 (Heinrich-Heine-University) was used for statistical power analysis. Differences between groups at p<0.05 (*) and p<0.01 (**) were considered as significant.

## 3. Results

### 3.1. Design of 2D and 3D PLA scaffolds for in vitro and in vivo experiments

In order to assess all aspects of the newly developed synthetic bone grafts from their *in vitro* biocompatibility to their efficiency to repair a critical-size bone defect in a large animal, the study was divided in four complementary parts (**Figure 1**): i) *in vitro* biocompatibility studies, notably cytotoxicity and stem cell response, on either film-coated or uncoated 2D PLA discs; ii) *in vivo* biocompatibility and biodegradability evaluation of 2D PLA discs in rat as small animal model; iii) assessment of the influence of scaffold geometry on BMP-2 loading in 3D mini-scaffolds; and iv) bone healing of critical-size metatarsal bone defect in sheep using BMP-2-containing 3D scaffolds of different geometries. **Figure 1** summarizes all experiments conducted and provides information on the custom-designed scaffolds, including their dimensions, experiment type, and qualitative and quantitative readouts.

First, *in vitro* biocompatibility assays were performed to ensure that PLA coated with the biomimetic film was not cytotoxic and favored a normal behavior of stem cells. 2D discs of different sizes that could be inserted in different types of multiwell-cell culture plates were fabricated to perform cellular assays according to NF EN ISO 10993 guidelines. Similar 2D discs with slightly different dimensions were used for subcutaneous implantations in rats to evaluate biocompatibility and biodegradability *in vivo* (**Figure 1**).

The external geometry of 3D porous scaffolds fabricated for the sheep study was designed based on the bone shape resected during osteotomy, i.e. a tube (cortical bone) with a central empty core (medullary cavity) (**Figure SI 1**). Therefore, cylinders of 25 mm in height and 14 mm in diameter with a differentiated inner ring were designed as external shape of the 3D scaffolds (**Figure 1**). Three types of geometric repeating unit cells (cubic, gyroid, and cubic-gyroid, a combination of cubic and gyroid) were selected for the inner structures according to the analysis of the literature. DOE were then performed to determine pore sizes as a function of the unit cell shapes. Three geometries were initially selected (**Figure 1**): i) Cubic S made of a thick hollow cylinder with open cubic pores of ∼0.87 mm and with a hollow inner ring of 5 mm in diameter. According to the predictive values obtained by the DOE, its mechanical properties were lower than those of sheep native bone, i.e. a compressive modulus of ∼300 MPa versus 28 GPa for native cortical bone and compressive strength of ∼7 MPa versus 16.6 MPa for native trabecular bone (no data was found for native cortical bone) [61,62]. ii) Gyroid L made of a thick hollow cylinder with ∼1 mm mean pore size and a hollow inner ring of 5 mm in diameter. Such structure was selected for its triply-periodic minimal surface design that is characterized by a zero mean curvature enhancing the surface area to volume ratio and that mimics trabecular bone structure [63]. According to the DOE, its compressive modulus was ∼100 MPa and its compressive strength ∼3 MPa. iii) Cubic-Gyroid made of an outer thick cylinder with ∼1.1 mm cubic opened pores and an inner ring of 7 mm in diameter filled with ∼2 mm gyroid pores. This latter geometry was selected as it combined the features of both cubic and gyroid structures, and we hypothesized that such composite structure may impact bone regeneration. According to the DOE, its compressive modulus and strength were respectively of ∼220 MPa and ∼6 MPa. Following the preliminary experiment in sheep, a fourth geometry was evaluated: iv) Gyroid S made of a thick hollow cylinder with ∼0.81 mm pores and a hollow inner ring of 5 mm in diameter. According to the DOE, its compressive modulus was ∼120 MPa and its compressive strength was ∼3.5 MPa. The porosity of all selected geometries was >75% according to the DOE.

In order to assess the impact of pore size and pore shape on the BMP-2 loading within 3D architectured scaffolds, 3D mini-scaffolds were specifically designed and printed (**Figure 1**). A geometry that was not considered for implantation was used in this BMP-2 incorporation study: Cubic L, a cubic geometry with pores of ∼1.3 mm. This geometry was used in the BMP-2 loading experiments since it had the same pore shape as Cubic S and the same surface as Gyroid L. Thus, it allowed to study the effect of pore size for two different pore shapes (cubic and gyroid) and pore shape for two different pore sizes (small (S) or large (L)) on BMP-2 loading. Cubic L was not considered for implantation in sheep metatarsal bone defect because it had too large pores, making it mechanically too brittle to be manipulated by surgeons.

It is worth noting that, in view of future clinical studies, clinical-grade (cgPLA) PLA was used for the fabrication of scaffolds tested *in vivo*, while *in vitro* experiments, except BMP-2 incorporation in 3D, were performed using regular-grade PLA (rgPLA) for financial reasons. Indeed, cgPLA costs ∼1000 times more than rgPLA. 2D discs and 3D scaffolds were designed and 3D-printed for the specific purpose of each experiment.

### 3.2. PLA scaffolds coated with a bioactive film are biocompatible in vitro

For direct cytotoxicity assay (**Figure 2a**), a material substrate is considered toxic if it promotes a reduction of 3T3 fibroblast viability by more than 30% compared to the one on negative control Thermanox substrate (set at 100%). Whereas cell viability was 41% on the positive control latex substrate, it remained higher than 94% on all PLA substrates tested, indicating that none of the biomaterial substrates induced cytotoxicity (**Figure 2a**). In addition, the viability of L929 cells in the presence of extracts from each of the 2D disc tested was always higher than 83% (**Figure 2b**), confirming the absence of cytotoxicity of all biomaterials tested.

**Figure 2.**
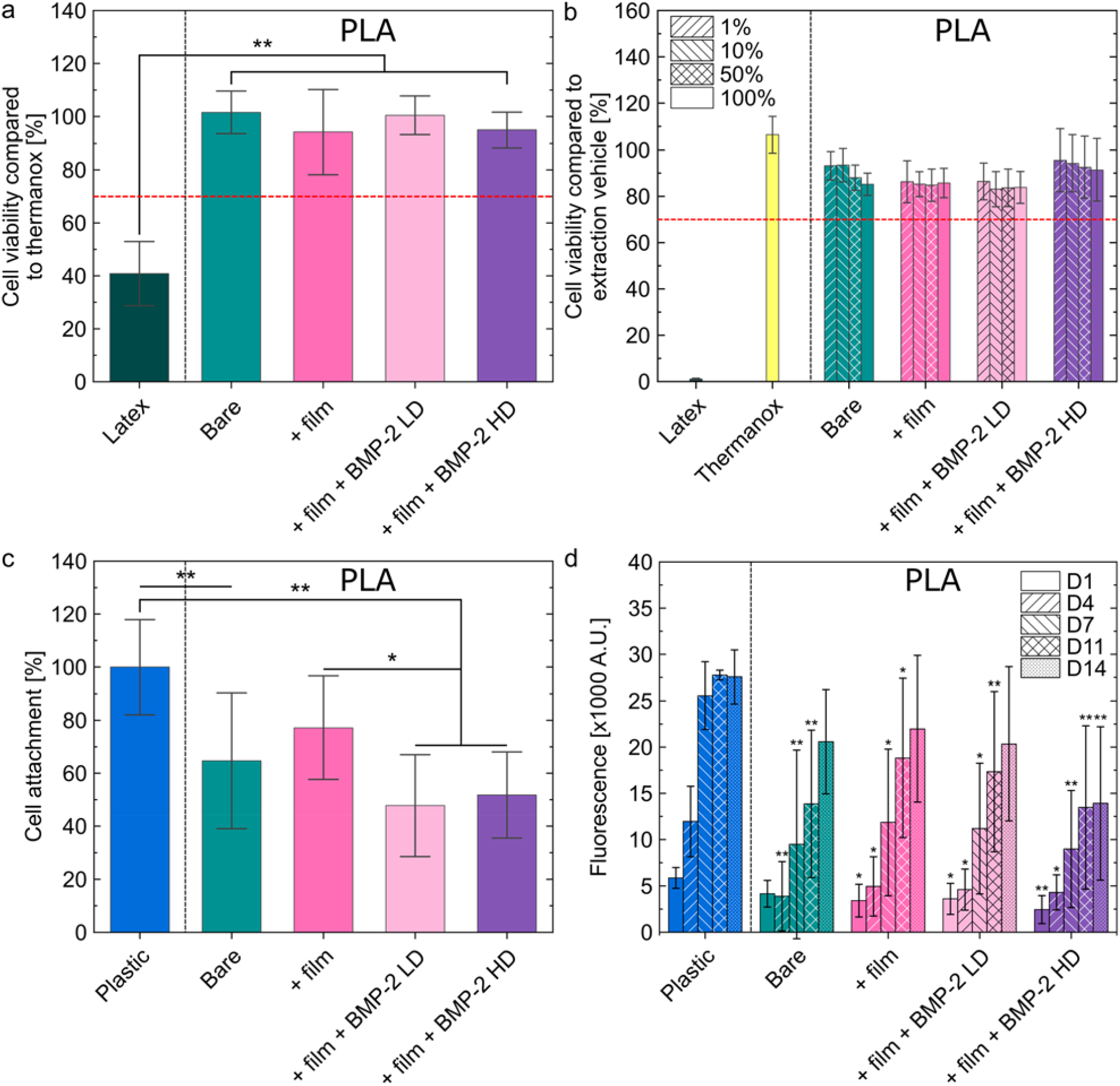
*In vitro* biocompatibility assays on 2D PLA discs. a) Direct cytotoxicity in contact. Cell viability compared to thermanox (expressed in %) is shown for each experimental condition: bare PLA discs, film-coated PLA discs and film-coated PLA discs loaded with two doses of BMP-2: LD and HD. b) Direct cytotoxicity with extract. Cell viability compared to an extraction vehicle (expressed in %) is shown for each experimental condition. c) Cell attachment (expressed as % in comparison to plastic) for each experimental condition. d) Cell proliferation, expressed as fluorescence arbitrary unit, for each experimental condition. Conditions were compared to the plastic control. Experiments were performed with *n* = 3 or 4 samples per condition in each independent experiment. *p<0.05; **p<0.01.

The analysis of hMSC attachment to biomaterial substrates after 15 h of culture showed that, compared to control plastic, ∼35% fewer cells attached onto the PLA surface in the absence or presence of the biomimetic film (**Figure 2c**). The presence of BMP-2 in the film further decreased cell attachment by ∼27%, independently of the BMP-2 dose. Additionally, the proliferative rate of hMSC was lower on PLA compared to control plastic substrate (**Figure 2d**). On plastic, cell number reached a plateau after around 7 days of culture, whereas no plateau was reached on bare PLA after 14 days of culture. However, a plateau was nearly reached on film-coated PLA with and without BMP-2 LD after 14 days, and a plateau was reached at 11 days on film-coated PLA loaded with BMP-2 HD. While coating PLA with the biomimetic film did not affect cell proliferation, the addition of BMP-2 decreased cell proliferation. This decrease was more marked with the highest BMP-2 dose. This may be explained by the fact that BMP-2 induced the osteoblastic differentiation of hMSC, thus blocking their proliferation. Differentiation tests should be conducted to assess this hypothesis.

Together, these results showed that 3D-printed PLA coated with the biomimetic film with or without BMP-2 was not cytotoxic. Furthermore, adding BMP-2 in the film reduced cell attachment but not in a dose-dependent way and increasing the BMP-2 dose decreased cell proliferation.

### 3.3. In vivo biocompatibility and biodegradability of film-coated PLA discs in small animals

To assess the biocompatibility and kinetics of resorption of film-coated PLA discs, 36 adults Wistar rats have been subcutaneously implanted with 4 samples each of either bare PLA, film-coated PLA without BMP-2, or film-coated PLA with BMP-2 loaded at two different BMP-2 concentrations: BMP-2 LD and BMP-2 HD (**Figure SI 2a**). The clinical and weight follow-up lasted 7, 28, or 48 days (*n* = 3 rats per time point, i.e. 12 samples per condition). Then, animals were euthanized and vital organs (liver, spleen, heart, brain, and kidney) and implanted tissue were explanted for histological analysis.

The clinical recovery was normal and healing of the skin evolved normally over time whatever the condition. The implant was always retrieved inserted into the skin, integrated into a fibrous capsule (**Figure SI 2b**). No infection occurred.

Histological analysis of the different vital organs did not reveal any histological abnormality (data not shown). Moreover, it was shown that the PLA discs never resorbed but were often not adherent to the surrounding tissue, with a fibrous capsule (yellow arrows) around them (**Figure 3a, e, and i**). A macrophagic inflammatory reaction with polynuclear cells (*) around the 2D PLA discs was observed at D7 (**Figure 3a’**), as expected for a normal reaction to a foreign body. This reaction decreased at D48 compared to D28 (**Figure 3e-l’**). The film (F) remained visible for all substrates tested at all time points (**Figure 3b, f, g, h, and l’**). However, it was sometimes peeled off from the PLA discs (**Figure 3c, d’, j, k’**) and some fragments were totally detached and surrounded by a macrophagic border with polynuclear cells (**Figure 3b’**) or giant multinucleated cells (#) (**Figure 3h’**), or they were encapsulated into a fibrous capsule (**Figure 3c’**) or in a vascularized fibroblastic shell (VFS) (**Figure 3j’**). Sometimes, macrophages were located next to the film (**Figure 3f’**). At D28 and D48, a vascularized fibroblastic shell in the periphery of the implant was observed (**Figure 3e’, f, i’, j, and k’**). This fibroblastic shell was delimited by a macrophagic border (red arrows) at D28 (**Figure 3e’**). No bone formation was observable for bare PLA and film-coated PLA without BMP-2 at any time points (**Figure 3a-b’, e-f’, and i-j’**). At D7, some new bone (NB) was forming in contact with the biomaterial with BMP-2 LD (**Figure 3c’**) and new bone was visible with BMP-2 HD (**Figure 3d**). At D28 and D48, new bone formation was observed for both BMP-2 doses (**Figure 3g-h’ and k-l’**), with sometimes bone tissue surrounded by osteoblasts (green arrows) (**Figure 3g’**). Histomorphometry showed that at D7, PLA + film + BMP-2 LD formed ∼1.5 times more bone compared to PLA + film + BMP-2 HD. On the contrary, at D28, 54% more bone was formed with BMP-2 HD compared to BMP-2 LD and at D48, 35% more bone was formed with BMP-2 HD compared to BMP-2 LD. The quantity of new bone formed with BMP-2 LD was not significantly higher at D28 and D48 compared to D7 because of high variability at D7. On the contrary, significantly more bone was formed at D28 and D48 compared to D7 when using BMP-2 HD. Interestingly, the quantity of newly formed bone did not increase between D28 and D48 (**Figure 3m**). A two-way ANOVA test was performed and showed that there was no significant difference in the quantity of new bone formed between BMP-2 LD and BMP-2 HD, but there was a significant difference between the quantity of bone formed at D7 and the quantity of bone formed at D28 and D48. The film present at the implant surface at each time point of the experiment was also evaluated: significantly more film was found to be present at D7 compared to D28 and D48, with however no significant difference between the experimental conditions.

**Figure 3.**
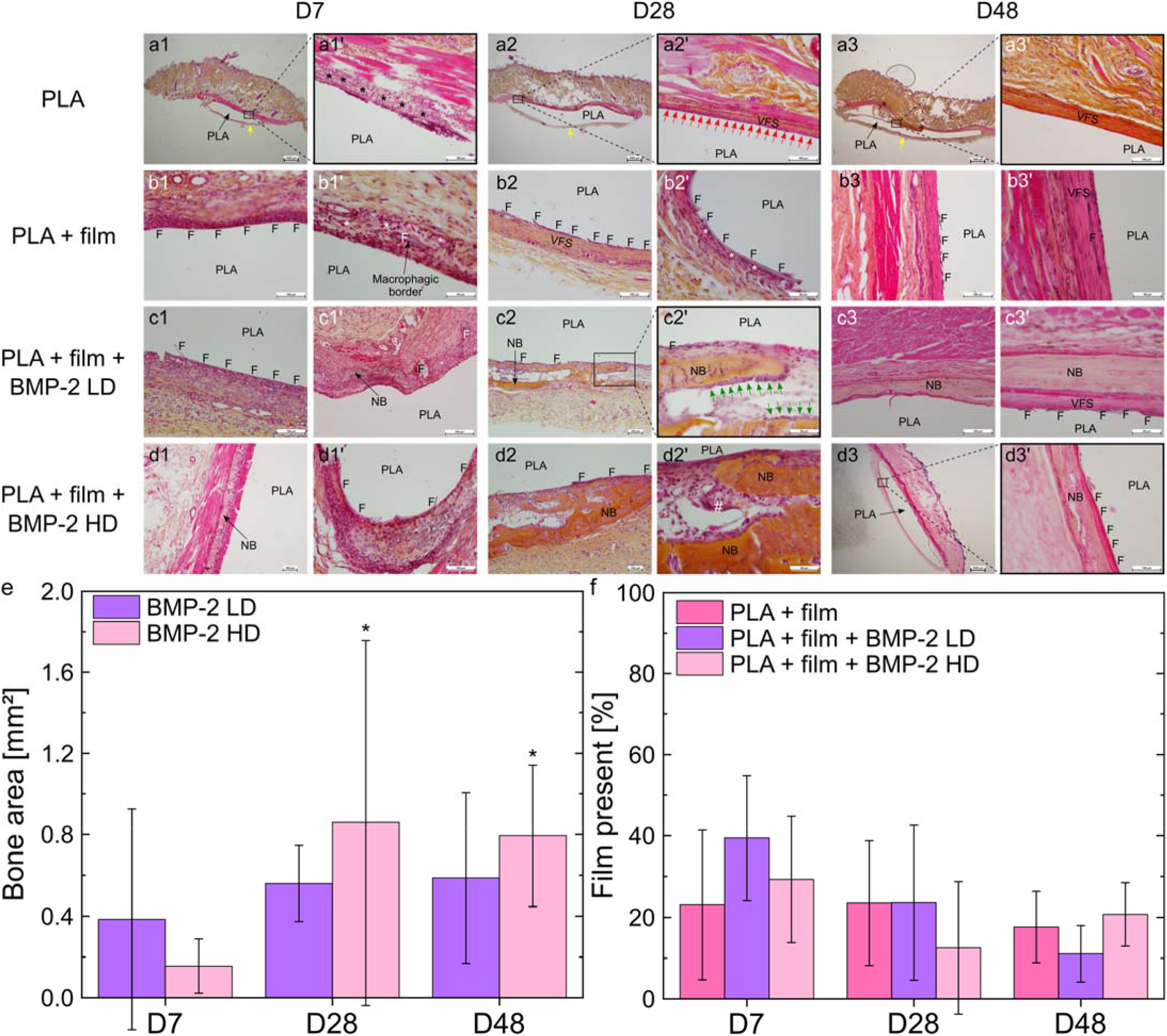
*In vivo* biocompatibility and biodegradability of films in rats quantified over time. Images of sections of PLA discs taken as a function of time at D7, D28, and D48 (endpoint) for the four experimental conditions studied: PLA; PLA + film; PLA+ film + BMP-2 LD; PLA + film + BMP-2 HD. a), e), i), and l) scale bar is 1 mm. a’), b), c)-d’), e’), f), g), h), i’), j), k), l), and l’) scale bar is 100 µm. b’), f’), g’), h’), j’), and k’) scale bar is 50 µm. Yellow arrows: fibrous capsule. (*) Macrophages and/or polynuclear cells. F: biomimetic film. NB: new bone. Red arrows: macrophagic border. VFS: vascularized fibroblastic shell. Green arrows: osteoblasts. (#) Giant multinucleated cells. m) Quantification of the bone area (mm^2^) formed at D28 and D48 when BMP-2 was used. * p<0.05 compared to D7. n) Quantitative analysis of the amount of film (%) remaining on the biomaterials surface. For each time point, there were *n* = 12 samples per condition using *n* = 3 rats, each receiving 4 implants.

### 3.4. BMP-2 incorporation in 3D scaffolds

The study of BMP-2 loading in the different 3D architectures was conducted *in vitro* in 3D mini-scaffolds made of cgPLA. µCT images of the different 3D mini-scaffolds, along with their surface, porosity, and pore size, are shown in **Figure SI 3**. The total amount of BMP-2 incorporated (in µg) is given for each type of geometry, as a function of BMP-2 concentration in the loading solution (**Figure 4a**). Effective surface doses incorporated (µg/cm^2^) are also given to take into account the differences in scaffold surfaces (**Figure 4b** and **Figure SI 4**). The % of BMP-2 from the initial loading solution effectively loaded in the scaffolds is also given (**Figure SI 5**). The incorporation of BMP-2 increased with the increase of BMP-2 concentration in the loading solution (**Figure 4a and b**). However, differences appeared between geometries: BMP-2 incorporation in Gyroid L appeared to increase linearly while it was not linear for the other geometries (**Table SI 2**). Moreover, BMP-2 loading in Gyroid S reached a plateau, while this was not the case for Cubic S and Cubic-Gyroid (**Figure 4a and b**). **Figure SI 4** shows that, at BMP10, Cubic L and Gyroid L incorporated significantly more BMP-2 than Cubic S and Gyroid S respectively, indicating that pore size influenced BMP-2 loading at low BMP-2 dose. At BMP30, Cubic L incorporated significantly more BMP-2 than Gyroid L, suggesting that pore shape influenced BMP-2 loading. At BMP50, Cubic S incorporated significantly more BMP-2 than Gyroid S, again suggesting an effect of pore shape on BMP-2 loading. Moreover, the BMP-2 incorporation in Gyroid S and Gyroid L were significantly different at BMP30 and BMP50. These results showed that pore size and shape, so scaffold geometry, influenced BMP-2 incorporation.

**Figure 4.**
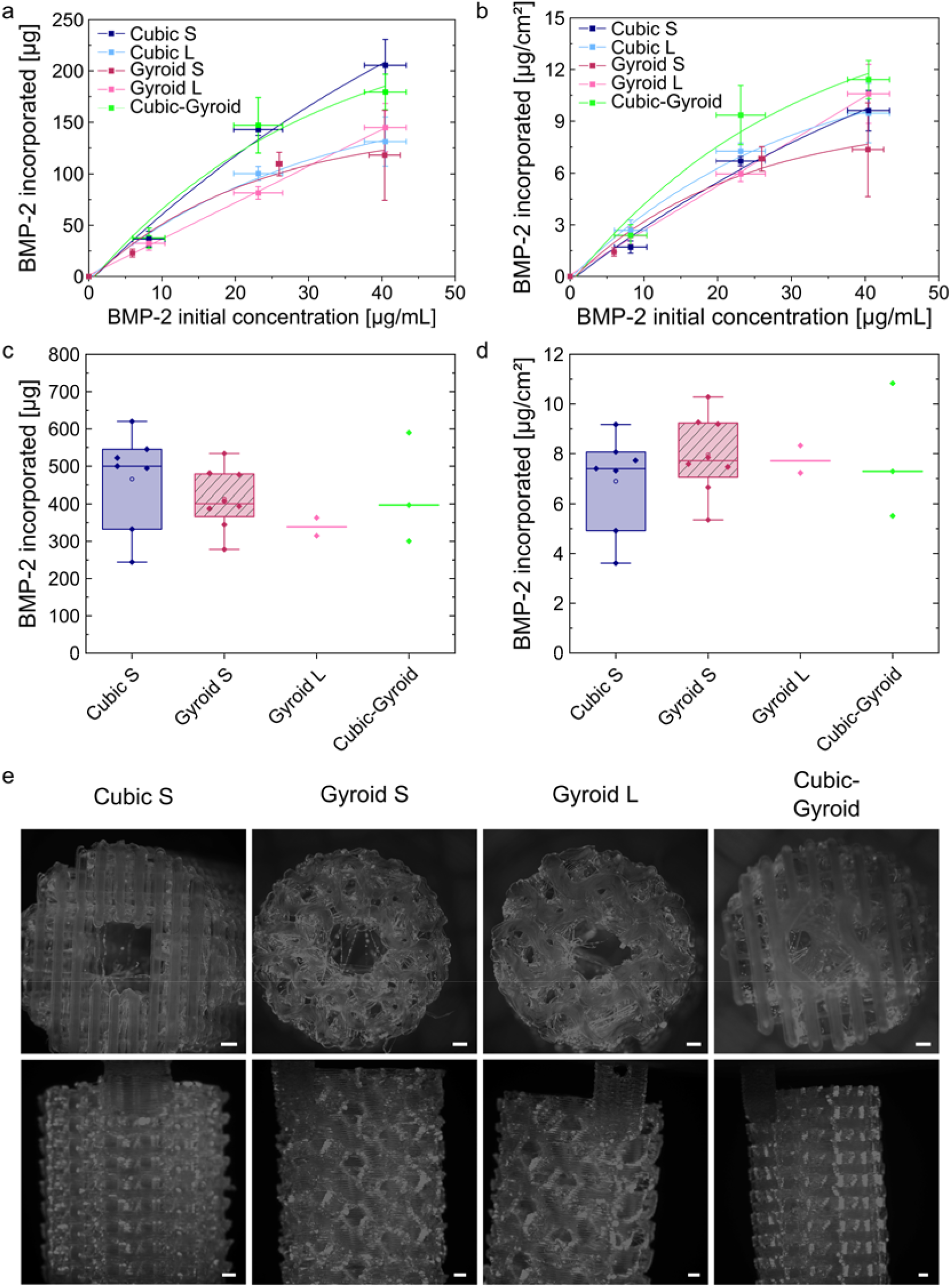
Characterization of BMP-2 loading in 3D mini-scaffolds. a) Quantification of the total dose of BMP-2 incorporated in 3D mini-scaffolds in function of the BMP-2 initial concentration in the loading solution expressed as absolute mass (µg) and b) expressed as surface dose in µg/cm^2^. The parameters extracted from the fits of the data are given in **Table SI 2a and 2b**, respectively. c) Total dose of BMP-2 incorporated in 3D scaffolds prepared for *in vivo* experiments in sheep (µg) as a function of scaffold geometry. d) BMP-2 dose incorporated in 3D scaffolds prepared for *in vivo* experiments in sheep expressed as surface dose (µg/cm^2^) after normalization by the scaffold effective surface, as a function of scaffold geometry. f) Fluorescence macroscopy images of 3D scaffolds loaded with BMP-2^Rhod^ for each studied geometry Cubic S, Gyroid S, Gyroid L, and Cubic-Gyroid. Scale bar is 1 mm.

For the *in vivo* implantation in a sheep metatarsal critical-size bone defect, a dose of 500 µg of BMP-2 per scaffold was targeted. As the scaffolds had different geometries, their surfaces were different and thus the concentration of BMP-2 in the loading solution was adapted to each scaffold geometry. The doses of BMP-2 incorporated into large 3D scaffolds used for *in vivo* implantation in sheep are shown in **Figure 4c and d**. There were some differences in BMP-2 incorporation between all geometries, but not statistically significant (**Figure 4c**): BMP-2 loading was the highest in Cubic S and the lowest in Gyroid L scaffolds. In terms of effective BMP-2 surface doses, similar BMP-2 doses were loaded in all geometries (**Figure 4d**).

Finally, the homogeneity of BMP-2 loading inside the 3D scaffolds was visualized by fluorescence macroscopy and microscopy using BMP-2^Rhod^ (**Figure 4e**, details in Supporting Information, and **Figure SI 6**). BMP-2 appeared to be homogeneously distributed within the 3D scaffolds. All together, these data showed that BMP-2 was efficiently loaded onto the surface of 3D porous scaffolds of various architectures designed for *in vivo* implantations, and that the incorporated BMP-2 dose was similar in all scaffolds.

### 3.5. Physico-chemical, mechanical, and morphological characterization of 3D PLA scaffolds and film coating prepared for in vivo implantation in sheep

Characterization of the different 3D scaffold geometries was performed using complementary techniques (**Figure 5**). First, the structural differences between the two types of PLA filaments, rgPLA and cgPLA, were analyzed using ATR-FTIR (**Figure 5a**) and SAXS (**Figure 5b**) to compare their chemical composition and crystallinity. ATR-FTIR showed no difference between the end-groups of both polymer chains, indicating that the filament structures were similar (**Figure 5a** and **Table SI 3**). Moreover, two peak ratios of interest, R_1_ (1209/1180 cm^-1^ band intensity) and R_2_ (1130/1080 cm^-1^ band intensity) indicate the degree of PLA crystallinity (the higher these ratios are, the more crystallized PLA is) [64]. Higher values were found for cgPLA (R_1_=0.62 and R_2_=0.67) compared to rgPLA (R_1_=0.54 and R_2_=0.52), suggesting that cgPLA was more crystallized than rgPLA (**Figure 5a**). Regarding SAXS, six diffraction peaks were identified for cgPLA compared to the three peaks identified for rgPLA, confirming the higher degree of crystallinity of cgPLA compared to rgPLA (**Figure 5b** and **Table SI 4**).

**Figure 5.**
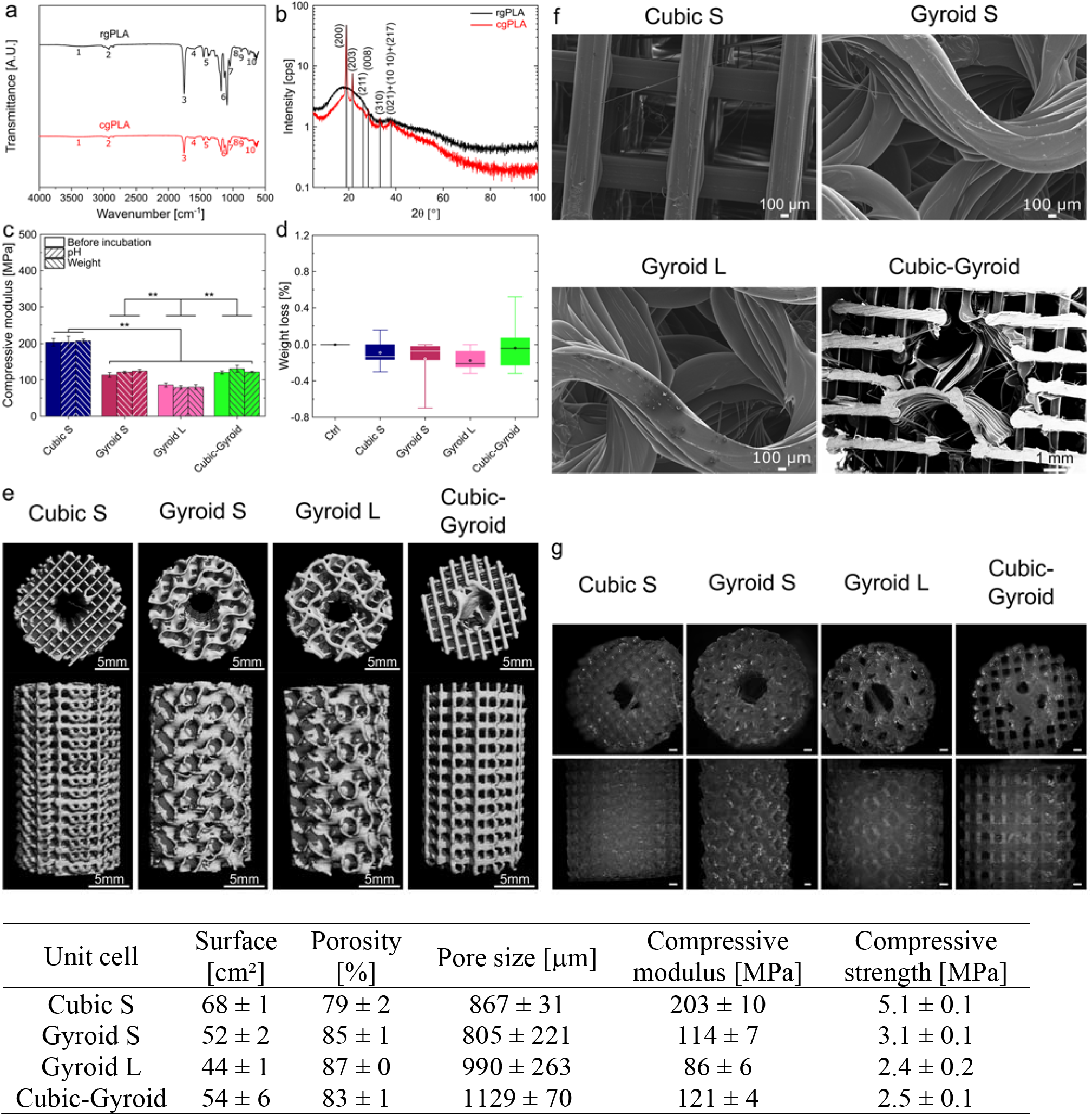
Physico-chemical, mechanical, and morphological characterization of PLA scaffolds and film coating. a) ATR-FTIR spectra (transmittance) of rgPLA and cgPLA. Remarkable peaks are numbered and identified in **Table SI 3**. b) SAXS spectra of rgPLA and cgPLA. Remarkable peaks are identified with Miller indices on the graph and in **Table SI 4**. c) Weight loss (expressed in %) measured for the scaffolds of different geometries Cubic S, Gyroid S, Gyroid L, and Cubic-Gyroid. d) Scaffolds’ compressive modulus (MPa) for the different scaffold geometries measured at different time points of the experiment: before incubation, after the immersion in a physiological solution over 12 weeks (scaffolds were never dried), and after the weight loss experiment, for which scaffolds were dried at each time point before weighting. e) µCT scans of the different scaffold geometries. f) Representative scanning electron microscopy images of the top surface of the different scaffolds for each geometry. g) Fluorescence macroscopy images of scaffolds coated with PLL^FITC^. Scale bar is 1 mm. h) Table recapitulating all measured values, including the effective surface (in cm^2^), porosity (in %), pore size (in µm), compressive modulus (in MPa), and compressive strength (MPa), for each scaffold geometry.

The *in vitro* degradation of the 3D PLA scaffolds was assessed by measuring the pH variation of the incubating solution (an indicator of the possible release of acidic products, **Figure SI 7a and b**), the scaffold weight loss (**Figure 5c** and **Figure SI 7c**), and the scaffold mechanical properties (**Figure 5d** and **Figure SI 7d**), which were measured in physiological conditions in a phosphate buffered saline solution (PBS) as a function of time (details in Supporting Information, **Figure SI 7**). The pH variation was the highest during the two first weeks of the incubation time and then decreased, but remained overall <4% for all scaffolds (**Figure SI 7a**). Gyroid L exhibited the lowest pH variation (**Figure SI 7b**). Cubic S with and without film coating, Gyroid S, and Cubic-Gyroid with film coating were different from the negative control, e.g. pure PBS without any scaffold. According to a two-way ANOVA statistical test with one factor being the geometry and the other one being the presence of the film, pH variation for Cubic S was statistically different from pH variation for Gyroid L, meaning that more acidic products were released by Cubic S (**Figure SI 7b**). The presence of the film on the scaffolds slightly increased pH variations for all tested geometries, indicating that more acidic products were released, although with minor changes (no statistical difference). The weight loss remained <1% over the incubation time, and was negative during the two first weeks, presumably due to water uptake by PLA fibers. Then, it progressively increased and was the lowest for Gyroid L (**Figure 5c** and **Figure SI 7c**). Mechanical properties of the 3D PLA scaffolds were assessed using uniaxial compressive tests (**Figure 5d** and **Figure SI 7d**). The compressive modulus and strength had lower values than those predicted by the DOE. Incubation in PBS during 12 weeks did not have a significant impact on scaffold mechanical properties as can be seen by the constant values of the compressive modulus and strength (**Figures 5d** and **Figure SI 7d**).

The 3D PLA scaffolds with different geometries were visualized with µCT scans (**Figure 5e**). Quantitative parameters were deduced from the µCT scans, including their effective total surface area (in µm^2^), porosity (given in %), and mechanical properties, which are summarized in **Figure 5h**. Cubic S and Gyroid L geometries displayed the highest (68 cm^2^) and the lowest (44 cm^2^) surfaces, respectively. Porosity values ranged from 79 to 87%, which should be sufficient to enable cell invasion and vascularization [65]. SEM was used to quantify pore sizes (**Figure 5f and h**), which were found to vary from 805 to 1130 µm. Cubic-Gyroid had the largest pores of ∼1130 µm and Gyroid S the lowest of ∼805 µm. Regarding the mechanical properties, the compressive modulus was in the range from ∼85 to 200 MPa and compressive strength from ∼2.5 to 5 MPa (**Figure 5h** and **Figure SI 8**). Cubic S had the highest compressive modulus (203 MPa) and compressive strength (5.1 MPa), while Gyroid L had the lowest (86 and 2.4 MPa respectively).

The homogeneity of the film coating onto the PLA fibers of porous 3D scaffolds was assessed by fluorescence macroscopy using PLL^FITC^ as last layer of the coating (**Figure 5g**) [66]. Film coating inside the bulk of the 3D scaffolds was also assessed after cutting them into pieces and imaging the slices. All geometries were fully and homogeneously coated by the film (details in Supporting Information and **Figure SI 9**). Film thickness in dry state was estimated to be ∼2 µm based on SEM imaging (details in Supporting Information and **Figure SI 10**).

### 3.6. Preliminary experiment to assess the influence of scaffold geometry on bone regeneration

A preliminary experiment with a reduced sample size (*n* = 2 per geometry) was conducted to select the optimal scaffold geometries (Cubic S, Gyroid L, or Cubic-Gyroid scaffolds). Scaffolds were implanted in 6 Pré-Alpes sheep, with one scaffold per sheep (**Figure SI 11**). Animals remained in good health. There were neither postoperative infection, nor implant failure. During explantation, it was impossible to distinguish macroscopically the newly formed bone from native bone or scar tissue around the implant.

The radiolucent property of the scaffold PLA material facilitated longitudinal X-rays analysis of the metatarsal bone defect. Time-dependent increase of the radiopacity throughout the defect was observed in both animals implanted with Cubic S, indicating an early and progressive bone formation in this specific scaffold. In contrast, a limited bone formation confined to the vicinity of the edges of the defect filled with either Gyroid L or Cubic-Gyroid scaffolds was observed up to 3 months post-implantation leading to a partial bone formation in these implants at 4 months (**Figure 6a**). One animal implanted with Cubic-Gyroid did not form bone at all (data not shown). These observations were confirmed by the X-ray scores, which increased over time and was the highest for Cubic S. **(Figure 6b and Figure SI 12)**.

**Figure 6.**
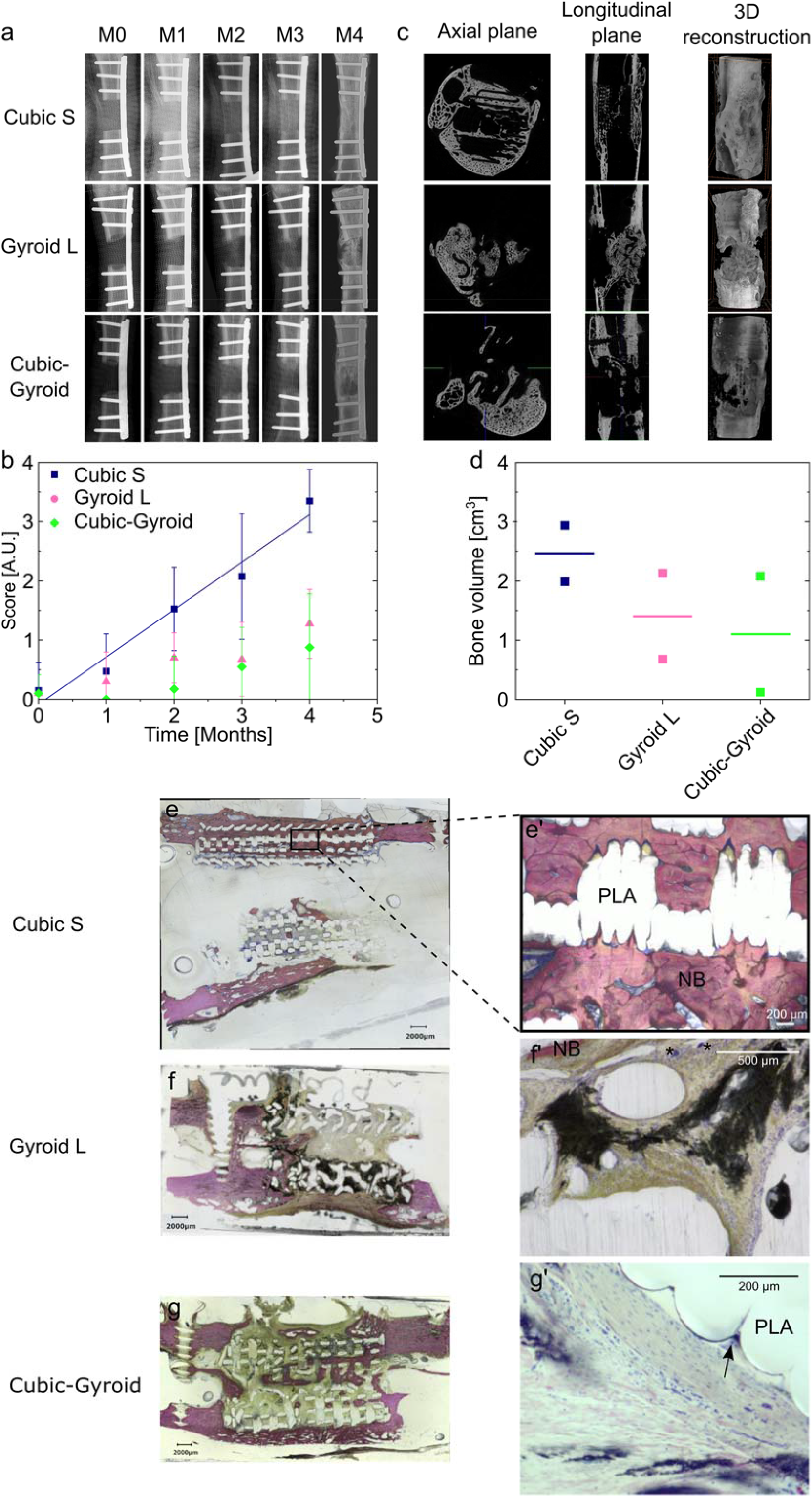
Preliminary experiment in sheep metatarsal critical-size bone defect to assess the influence of scaffold geometry on bone regeneration. a) Representative X-ray scans of bone regeneration achieved with film-coated scaffolds loaded with BMP-2 with different internal geometries acquired at different time points: right after scaffold implantation (M0), after one, two, three months (M1, M2, M3), and after explantation (M4). b) X-ray score given by the clinician as a function of the time for each scaffold geometry. Data are represented as mean ± SD of scores given by 5 clinicians and veterinarians. The scores for Cubic S were linearly fitted (R^2^ = 0.97). c) Representative µCT scans acquired after explantation for each scaffold geometry. For each geometry, scans in the axial plane, longitudinal plane, and a 3D reconstruction are shown. d) Quantification of the newly formed bone volume for each scaffold geometry. e)-g’) Representative histological sections for each scaffold geometry: Cubic S, Gyroid L, and Cubic-Gyroid. For each geometry, a global section and a magnified view are given. Bone is stained in pink. NB: new bone. (*) Giant cells. Black arrow: multinucleated giant cells.

µCT scans showed that the new bone grew in Cubic S primarily in the continuity of the cortices leading to bone union, whereas it grew non-uniformly in the defects filled with Gyroid L and Cubic-Gyroid scaffolds (**Figure 6c**). Bone quantification confirmed that Cubic S geometry promoted higher bone formation compared to the other geometries (mean of 2.5 cm^3^, **Figure 6d**). Bone homogeneity was also higher with Cubic S (**Figure SI13 and Figure SI14**). Interestingly, the newly formed bone in Cubic S was similarly distributed in all areas of the defect, namely proximal, central and distal bone while it tended to be more localized in the proximal and distal areas in Gyroid L and in the proximal area in Cubic-Gyroid. Ectopic bone formation was also lower in Cubic S than with the other geometries (**Figure SI13 and Figure SI14**).

The histological examination indicated abundant new bone tissue in the pores and around pores of Cubic S scaffolds (**Figure 6e**). Scaffold struts were either surrounded by newly formed bone (NB) (**Figure 6e’**) or by a fibrous tissue. More fibrous tissue was observed within the Gyroid L and Cubic-Gyroid scaffolds with the presence of islets of new bone (**Figure 6f and g**) but a larger magnification of another section shows some inflammatory cells and giant cells (*) (**Figure 6f’**). Cubic-Gyroid led to some new bone formation inside the scaffold (**Figure 6g**), and some multinucleated giant cells were observed (**Figure 6g’**, black arrow). Globally, this histological analysis confirmed scaffold geometry influenced bone regeneration and that Cubic S led to the best bone regeneration among the three geometries tested.

Based on this preliminary experiment, we decided to pursue our investigations with Cubic S, which induced the highest amount of newly formed bone, had high mechanical properties and high porosity. For the second main experiment, we decided to add a gyroid geometry with a pore size similar to Cubic S, i.e. ∼805 µm, hereafter named Gyroid S. These two geometries only differ in their pore shape.

### 3.7. Main experiment to study the influence of pore shape and optimize bone regeneration

BMP-2-loaded Cubic S (*n* = 5, completed with the 2 sheep from the preliminary experiment to reach a sample size of *n* = 7) and Gyroid S (*n* = 7) scaffolds as well as 2 controls per geometry (film-coated scaffolds without BMP-2) were implanted in the sheeps. Simimarly to the preliminary experiment, animals remained in good health with no postoperative problem, and the newly formed bone was well integrated.

X-ray scans and X-ray scores were acquired and analyzed respectively as in the preliminary experiment (**Figure 7a and b**). Limited bone formation was formed to the vicinity of the cut edges of the defects in both types of scaffolds implanted without BMP-2 (**Figure 7a**). Full bridging of the defect was observed in 6/7 animals implanted with Cubic S + BMP-2 (**Figure 7a**), whereas 1/7 animal led to partial bridging of the defect (**Figure SI 15a**). When implanted with Gyroid S + BMP-2, full bridging of the bone defect been achieved in 3/7 animals (**Figure 7a**), partial bridging in 1/7 animal (**Figure SI 15b**), and scarce bone formation in 3/7 animals (**Figure SI 15c**). These observations were confirmed by the X-ray scores, which steadily increased with the implantation time in animals implanted with both types of scaffolds containing BMP-2. As expected, scaffolds loaded with BMP-2 displayed significantly higher X-ray scores and indexes compared to scaffolds without BMP-2 (**Figures SI 16 and SI 17**). The defects filled with Cubic S + BMP-2 showed significantly higher X-ray scores at all time points, except at 2 months, compared to Gyroid S + BMP-2 (**Figure 7b** and **Figure SI 16**). The gap between both geometries increased with time. X-ray scores were fitted with an exponential function providing a plateau value B_max_ which was found to be slightly lower for Cubic S + BMP-2 than for Gyroid S + BMP-2 (5.3 vs. 5.6). However, this plateau was reached faster for Cubic S + BMP-2 scaffolds (τ value of 5.4 months vs. 9.3 months, respectively for Gyroid) (**Figure 7b**). Defects implanted with Cubic S + BMP-2 showed significantly higher bone filling and homogeneity indexes and lower ectopic bone formation index than defects with Gyroid S + BMP-2 **(Figure SI 17)**.

**Figure 7.**
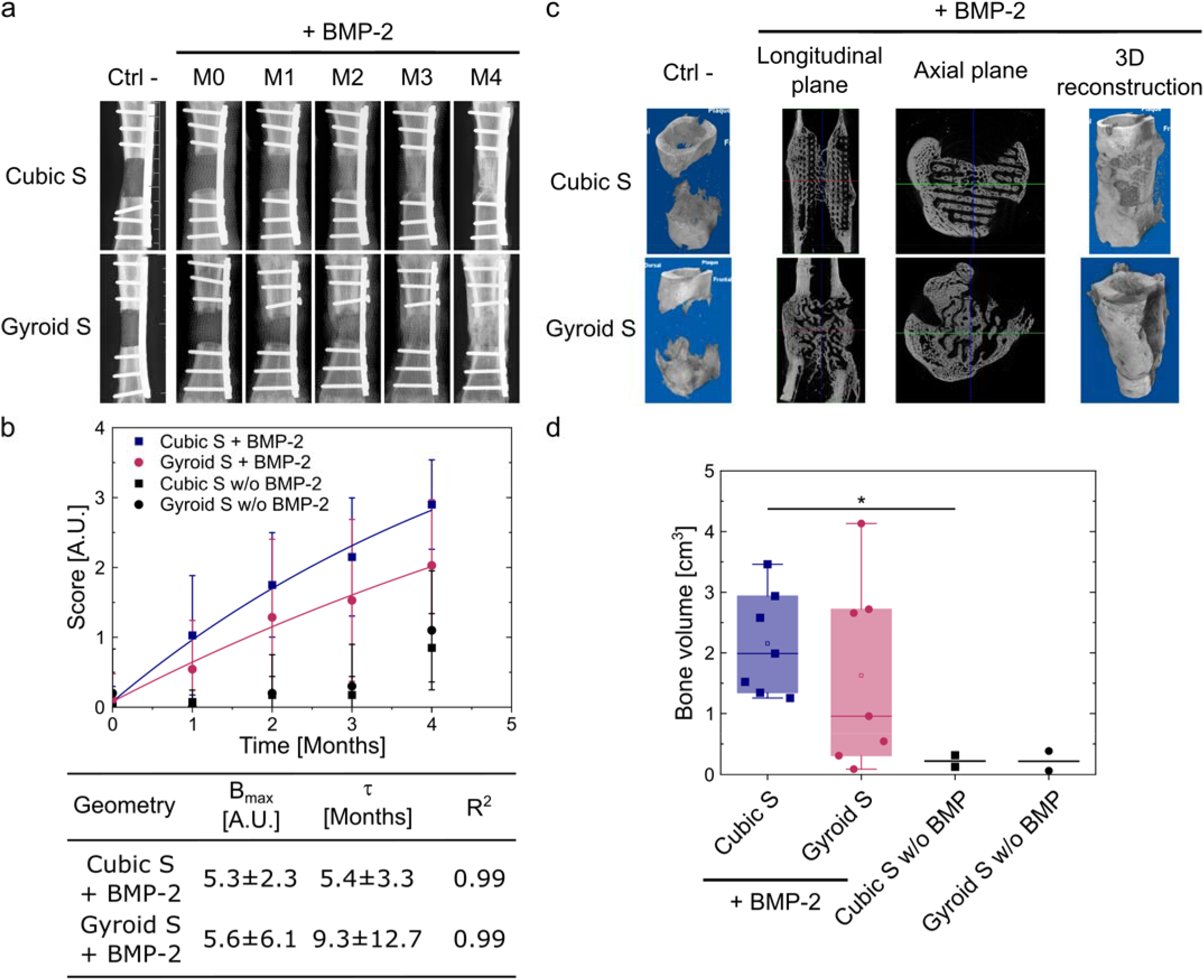
Main experiment in sheep metatarsal critical-size bone defect to assess the influence of pore shape and optimize bone regeneration. a) Representative X-ray scans acquired at different time points: right after scaffold implantation (M0), after one, two, three months (M1, M2, M3), and after explantation (M4) for each scaffold geometry. b) X-ray score as a function of the time (same calculation as in **Figure 6**). The scores for Cubic S and Gyroid S loaded with BMP-2 were fitted with an exponential function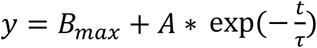. Quantitative parameters extracted from the fits are given (B_max_ and τ) in the corresponding table. d) Quantification of the newly formed bone volume for each scaffold geometry.

µCT scans provided evidence that a clinical bone union (defined as ≥ 3/4 united cortices) occurred in 5/7 and 3/7 animals implanted with Cubic S + BMP-2 and Gyroid S + BMP-2, respectively. This observation differed from the one made from X-rays for Cubic S + BMP-2 scaffolds on which 6/7 animals led to bone union. This difference between these two observations comes from the more limited field of view provided by 2D X-rays compared with 3D µCT scans. Quantitatively, Cubic S + BMP-2 induced higher bone formation than Gyroid S + BMP-2 (mean of 2.2 cm^3^ vs. 1.6 cm^3^, **Figure 7d**) but the difference was not statistically significant (p=0.38). Additionally, the bone volume was significantly different in Cubic S + BMP-2 compared to Cubic S w/o BMP-2. However, it was not different in Gyroid S + BMP-2 compared to Gyroid S w/o BMP-2 presumably due to the highly variable bone volume outcomes obtained with Gyroid S + BMP-2 (**Figure 7d**). In the presence of BMP-2, the newly formed bone was homogeneously distributed within the scaffold at proximal, central and distal locations, for both Cubic S and Gyroid S geometries. In striking contrast, in the absence of BMP-2, the new bone was mainly formed in the proximal area (**Figure SI 18**).

The histological analysis of the tissue sections of defects showed that, in the absence of BMP-2, the new bone deposition within the implant was scarce, confined to the bone ends of the defect, localized either at the periphery or in the core of the PLA scaffold, but never inside the scaffold pores ((**Figure 8a** with Cubic S, **Figure 8b** with Gyroid S and **Figure SI 20**). These observations indicated that the film-coated PLA scaffold was not osteoconductive by itself. Bone distribution inside the BMP-2-containing scaffolds showed some variability between animals (**Figures 8c, 8c’, 8d and 8d** and **Figures SI 19 and SI 20**). In both groups, some areas of the scaffolds were filled with bone while others remained empty, providing evidence of a nonuniform bone induction. In addition, in most of the explants, the new bone tissue was mostly formed in the outer part or outside of the PLA scaffolds (peripheral area) in the continuity of the cortices, leading to a new connecting cortex when the material implant was not aligned with the native cortices (**Figures 8c, 8c’, 8d and 8d and Figure SI 19c, c’, e, and e’**). At higher magnification, the bone tissue formed around the scaffolds appeared mature (lamellar), homogeneous and dense with thick trabeculae filled with bone marrow **(Figures 8e and 8g)**. When found within the scaffold pores, the new bone tissue (NB) displayed either lamellar features with the presence of blood vessels (V) **(Figure 8f)** or woven bone features with osteocytes, bone-lining cells and osteoblasts depositing osteoid tissue, thus, revealing an active bone formation (**Figures 8i and 8j)**. However, we noted that the new bone was scarcely in contact with the PLA material struts **(Figures 8g, 8i, 8j)**. Isolated bone nodules were also observed at a distance from the PLA struts (**Figure 8g**). Besides these bone nodules, infiltrated tissue including fibrous connective tissue containing blood vessels was present **(Figures 8g, 8i, 8j)**. Numerous multinucleated giant cells (red arrowheads) were found close to or in contact with the implant PLA material (**Figures 8i and 8j)**. Some remnants of the biomimetic film (F) were also observed close to the scaffold’s struts, encapsulated into a fibrous capsule and most often surrounded by multinucleated giant cells (**Figures 8i and 8j**).

**Figure 8.**
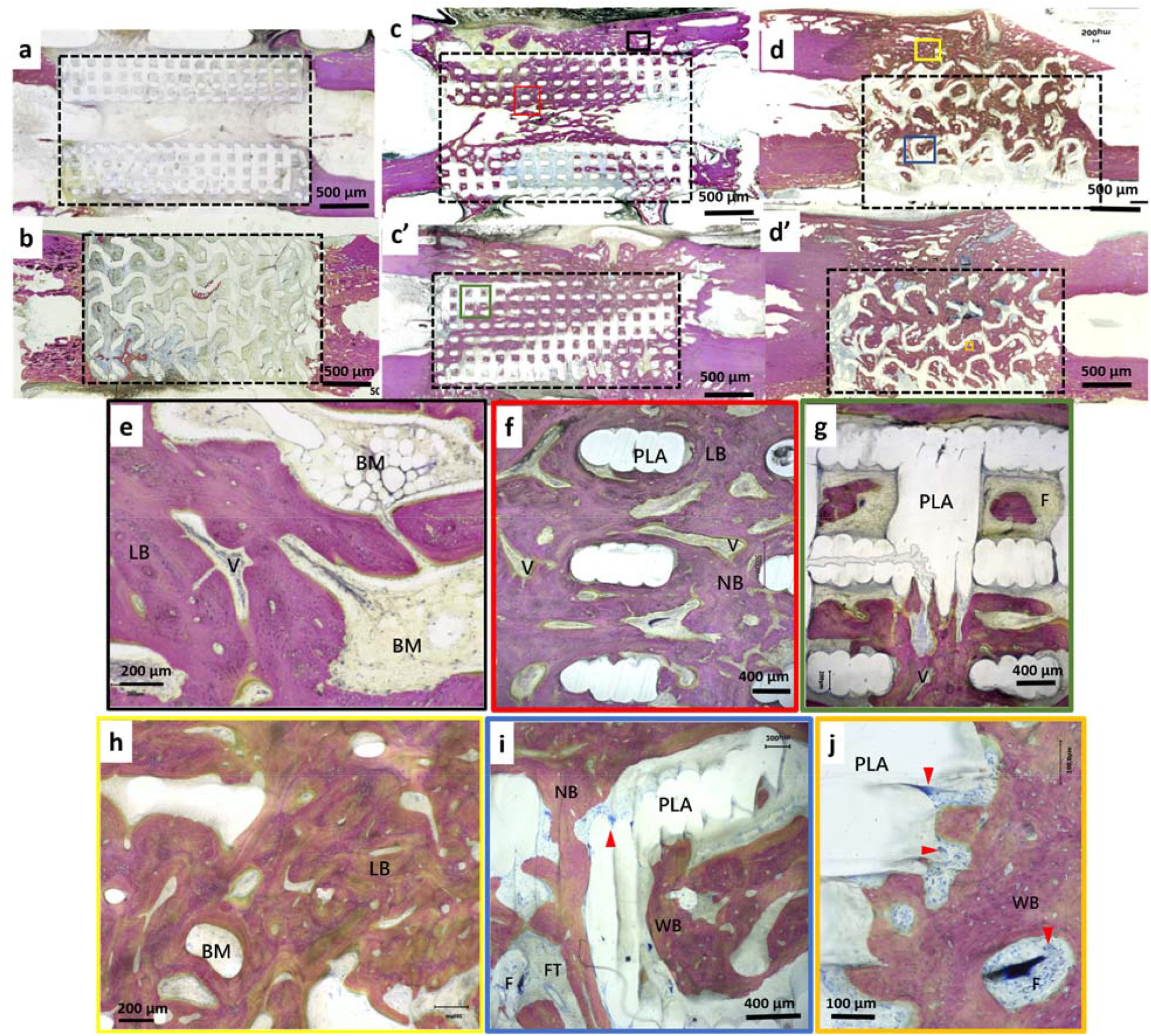
Histological analysis of scaffolds with Cubic S and Gyroid S geometries. Representative histological sections the different types of scaffolds. Film-coated PLA scaffold without BMP-2 a) Cubic S and b) Gyroid S geometries. In the presence of BMP-2 in the films, bone was formed in both types of scaffolds c-c’) Cubic S and d-d’) Gyroid S. At higher magnification, specific features of the newly formed bone were visible : e,f) formation of mature lamelar bone (LB), homogeneous and dense bone with thick trabeculae filled with bone marrow (BM). Vessels (V) are also visible. g) some remnants of the films (F) are visible, encapsulated in a fibrous capsule. h) lamelar bone and bone marrow; i) Numerous multinucleated giant cells (red arrowheads); j) these cells were visible close to the scaffold struts. Bone is stained in pink. PLA: PLA scaffold. BM: bone marrow; F: biomimetic film; LB: lamellar bone; NB: new bone. WB: woven bone. V: blood vessels. Scale bars: a-d: Scale bar is 500 µm. f,g,i: 400 µm. e,h: 200 µm. j: 100 µm.

In summary, this main experiment provided evidence that BMP-2 containing scaffolds consistently induced bone tissue either inside the 3D architectured scaffolds and/or at the periphery of the scaffold, while only a very limited amount of bone tissue was found in and around the scaffolds without BMP-2. Another finding of this experiment is that Cubic S + BMP-2 scaffolds led to a higher amount of newly formed bone when compared with Gyroid S + BMP-2 scaffolds. The variability of the newly formed bone volumes was lower and the kinetics of bone formation was faster for Cubic S + BMP-2 compared to Gyroid S + BMP-2.

## 4. Discussion

In this study, we used the FDM technique to design and fabricate PLA scaffolds specifically for different *in vitro* and *in vivo* assays, showing the high versatility of this 3D printing technique. The characterization of PLA of clinical grade, a material required for clinical translation, revealed specific differences with PLA of regular grade, notably in terms of crystallinity. This observation highlights the need to perform the experiments with the appropriate raw material.

Results from the biocompatibility experiments (**Figure 2**) showed that hMSC adhesion was reduced on BMP-2-loaded films and cell adhesion was similar for BMP-2 loaded at 30 versus 60 µg/mL, indicating that a plateau was probably reached. This appears to be in contradiction with previous results obtained on C2C12 cell adhesion [34] and on human periosteum derived stem cells (hPDSCs) [71]. However, this effect may be due to the BMP-2 doses used here, respectively 30 µg/mL and 60 µg/mL, which were rather high for *in vitro* studies. Indeed, Sales et al. [71] found that BMP-2 concentrations above 5 µg/mL led to a decrease in hPDSCs adhesion. For C2C12 cells, a plateau was reached from 2.5 to 20 µg/mL. Thus, there may be dose-dependent responses to BMP-2 depending on the cell type.

Preclinical studies are a prerequisite to develop bone tissue engineering products, and safety and efficiency should be proved in relevant animal models. Critical-size bone defects are of particular relevance in view of their close resemblance to clinical situations. Sheep is a particularly good candidate to study bone regeneration in orthopedics because its long bones are of similar size compared to humans and its bone physiology, biomechanics, stem cells, and response to BMP-2 are comparable to that in humans [1,54– 56,67,68]. Compared to other long bones such as tibia and femur, metatarsus displays a more hostile environment for bone regeneration as it is encircled by tendons and lacks muscle coverage, a major source of periosteal vascularization, progenitor cells and paracrine stimuli [69]. In addition, previous studies from members of our team showed that consistent bone union was achieved in that model when defects were filled with bone autografts, the current standard of care in clinical situations [52,60]. Thus, the sheep metatarsal bone appeared as an appropriate model to assess a newly developed bone graft substitute. A similar bone graft substitute made by 3D printing of PLA already proved to be efficient in the more favorable environment which was the minipig mandible [53]. Indeed, though the presence of teeth and proximity with saliva induces unfavorable bacteriological conditions, the minipig mandible displays a large amount of well-vascularized cortico-cancellous bone with extensive muscle coverage, which creates more conducive conditions for bone regeneration [70]. Here, we proved that the bone graft substitute could also regenerate bone in a hostile and load-bearing environment.

Controlling the BMP-2 dose is of prime importance in the clinical applications, since it was already shown that a too high BMP-2 dose can lead to adverse effects [72]. As a result, the BMP-2 dose should be carefully adapted to ensure an optimal bone regeneration without side effects. In this study, a BMP-2 dose of ∼80 µg/cm^3^ of defect was targeted, based on our previous study on minipig mandible [53] where an efficient bone regeneration was found for this dose. BMP-2 incorporation in the 3D scaffolds was found to be higher in the present study compared to the previous one, with BMP-2 loaded dose of ∼120 µg/cm^3^ of defect. In previous studies in sheep, Yang et al. used a BMP-2 dose of 400 µg/cm^3^ of defect to repair a 5 cm-long tibial defect [73]. According to them, studies in preclinical large animal models use doses from 100 to 800 µg/cm^3^ (in Yang: [72,74–76]). In another study, Maus et al. repaired a trepanation defect in sheep distal femoral epiphysis with a BMP-2 dose of 200 µg/cm^3^ of defect [77]. Similarly, Cipitria et al. repaired a sheep metatarsal critical-size bone defect using a BMP-2 dose of ∼190 µg/cm^3^ of defect [56]. Thus, in the other studies, the BMP-2 doses were between 1.6 and 4-fold higher than in the present one. In addition, a dose of ∼120 µg/cm^3^ represents a 12-fold decrease compared to 1.5 mg/mL loaded into collagen sponges used in clinics. Thus, this low BMP-2 dose delivered via the surface coating of a 3D-printed scaffold enabled an efficient and safe bone regeneration without adverse effects in our model.

In our previous study on minipig mandibles [53], we developed a CT-scan score to qualitatively assess bone regeneration over time. Here, a similar score was used to qualitatively assess bone formation based on X-rays (**Figures 6b and 7b**), the initial score being slightly adapted by considering only one type of bone, without distinction between cortical and cancellous bones, which can barely be distinguished on radiographs. We found that this simple score, which can be given by clinicians, agrees well with the quantitative results obtained by analyzing µCT images (**Figure SI 21**). This finding is of great interest, since it provides an easy and fast qualitative assessment of the amount of newly formed bone. It can be a timesaving predictive tool of the synthetic graft (peut on parler de greffe pour un matériau de synthèse ?) success, without the need to wait for µCT images.

In this study, we compared the efficiency of different scaffold geometries to repair a critical-size sheep metatarsal bone defect. We found that a cubic geometry with a mean pore size of ∼870 µm could efficiently regenerate bone in 5/7 cases. Comparatively, the gyroid geometries tested with mean pore sizes of ∼1 mm and ∼805 µm did not regenerate bone so consistently (for Gyroid L, 2/2 scaffolds partially bridged the bone defect and for Gyroid S, 3/7 fully bridged the defect, 1/7 partially bridged it, and 3/7 did not bridge it at all). These results are contrasting with those of found by Van Hede et al., who compared cubic (called orthogonal in their study) and gyroid structures in a calvarium rat model with a pore size of 700 µm [50]. The different results may be explained by the different implantation sites (metatarsal bone vs. calvarium), the animal model used (sheep vs. rat), the scaffold material (polymer vs. ceramics), and the presence or absence of BMP-2 (no BMP-2 was added in Van Hede et al. study). Notably, the implantation in Van Hede et al. was subperiosteal (extremely favorable environment) without any real loss of substance, especially without any critical-size bone defect [50]. Another reason may be that Cubic S has more opened pores located at the top and bottom sides of the scaffolds, which were in contact with bone edges, compared to Gyroid S. These opened pores may promote fluid infiltration inside the scaffolds, so that bone precursor cells better penetrate in the core of the scaffold. This was the opposite in the study of Van Hede et al. where gyroid scaffolds had more opened pores in contact with bone compared to scaffolds with orthogonal pores. Besides pore size and shape, another parameter has been identified by researchers as important for bone regeneration: permeability [78–80]. The ratio of scaffold surface over volume may also play a role [81,82] since the higher the ratio will be, the more surface will be available for cells to adhere. When considering the volume of the defect, surface/volume ratio was the highest for Cubic S (1.8) followed by Cubic-Gyroid (1.4), Gyroid S (1.35), and Gyroid L (1.1).

This study presents some limitations. Indeed, the number of animals used may be considered low. However, it is difficult to propose studies with larger groups sizes when working with large animals and it is desired to limit the number of animals used to respect the 3Rs principle. A priori statistical power analysis showed that for an effect size set to 0.5, α set at 0.05, and desired power set at 0.80, 64 sheep would be needed per group to see a statistical difference, which is inconceivable. If keeping the groups size at 7, the effect size should be > 1.6 in order to reach a power > 0.80. A post-hoc statistical power analysis showed that power was 0.11 when comparing Cubic S + BMP-2 vs. Gyroid S + BMP-2 (effect size of 0.4); 0.92 for Cubic S + BMP-2 vs Cubic S w/o BMP-2 (effect size of 3.2); and 0.28 for Gyroid S + BMP-2 vs Gyroid S w/o BMP-2 (effect size of 1.3). Another limitation is the incomplete loading of implants during the study due to the cast and walking bar maintaining the limb. This incomplete loading may have led to less bone formation since it is known that mechanical constraints are required for efficient bone repair. Furthermore, we noticed that when the diameter of the scaffold was smaller than the diameter of host bone, it led to a misalignment of the scaffold with bone and thus to less bone formation. Ideally, the scaffold length should be perfectly adapted to the bone defect to avoid delayed or poor osseointegration [83]. This highlights the necessity to fabricate personalized scaffolds that would be perfectly adapted to the specific defect in each animal.

## 5. Conclusion

We designed PLA scaffolds with different internal geometries, fabricated them by FDM, and coated them with a biomimetic polyelectrolyte film delivering BMP-2 to repair a critical-size metatarsal bone defect in sheep. Film-coated PLA loaded with BMP-2 was proved to be biocompatible both *in vitro* and *in vivo*. By tuning scaffold internal geometry, we showed that scaffold geometry influenced BMP-2 incorporation and bone regeneration. X-ray scans, µCT scans, and histology proved that scaffolds with cubic pores of ∼870 µm loaded with BMP-2 at ∼120 µg/cm^3^ led to the formation of new bone without any adverse effects. The new bone formed homogeneously in the longitudinal direction of the bone defect. Notably, the BMP-2 dose used here was ∼12 fold lower than in the commercially available collagen sponges, and ∼1.6 to 4 fold lower than comparative studies in large animals. Furthermore, the clinical score given by clinicians on the X-ray scans acquired during animal follow-up revealed to be an easy predictive tool of the quantitative assessment of bone volume done by µCT scans. This work opens perspectives for a future personalized treatment of large bone defects in patients, by adapting scaffold shape and size for each patient and precisely controlling the BMP-2 dose delivered via the film coating of the 3D scaffold.

## CRediT author contribution statement

**CG:** Conceptualization, Methodology, Software, Validation, Formal analysis, Investigation, Resources, Data curation, Writing – original draft, Writing – review and editing, Visualization. **SC:** Formal analysis, Investigation, Data curation, Visualization. **CM:** Conceptualization, Methodology, Validation, Formal analysis, Investigation. **PM:** Conceptualization, Methodology, Software, Validation, Investigation, Resources. **CD:** Conceptualization, Methodology, Validation, Formal analysis, Investigation. **SR:** Conceptualization, Methodology, Validation, Formal analysis, Investigation. **JVial:** Investigation. **JVollaire:** Methodology, Validation, Investigation, Resources. **MR:** Conceptualization, Methodology, Validation, Formal analysis, Investigation. **HG:** Investigation, Resources. **HE-H:** Methodology, Investigation, Data curation. **AC:** Investigation. **VJ:** Validation, Resources, Writing – review and editing, Supervision, Project administration, Funding acquisition. **LB:** Supervision. **GB:** Conceptualization, Methodology, Validation, Formal analysis, Resources, Writing – review and editing, Supervision, Funding acquisition. **MD:** Writing – original draft, Writing – review and editing, Supervision, Project administration, Funding acquisition. **MM:** Conceptualization, Methodology, Validation, Formal analysis, Investigation, Resources, Data curation, Supervision, Project administration, Funding acquisition. **VV:** Conceptualization, Methodology, Validation, Formal analysis, Investigation, Resources, Data curation, Writing – review & editing, Supervision, Funding acquisition. **DL-A:** Conceptualization, Methodology, Validation, Formal analysis, Resources, Data curation, Writing – original draft, Writing – review & editing, Visualization, Supervision, Project administration, Funding acquisition. **CP:** Conceptualization, Methodology, Validation, Writing – original draft, Writing – review and editing, Supervision, Project administration, Funding acquisition.

## Data availability statement

The data supporting the findings of this study are available upon request to the corresponding authors.

## Declaration of competing interest

The authors declare that they have no known competing financial interests or personal relationships that could have appeared to influence the work reported in this paper.

## Acknowledgments

This work was supported by the Agence Nationale de la Recherche (ANR-18-CE17-0016, OBOE), the Fondation “Gueules Cassées” (contract N°33-2022), and by the European commission European Research Council (ERC BIOMIM GA259370, ERC PoC BioactiveCoatings GA692924 and ERC PoC REGENERBONE GA790435). CP is a senior member of the Institut Universitaire de France whose support is also acknowledged. The Optimal imaging platform is supported by France Life Imaging (French program “Investissement d’Avenir” grant; “Infrastructure d’avenir en Biologie Santé”, ANR-11-INBS-0006) and the IBISA French consortium “Infrastructures en Biologie Santé et Agronomie”.

The authors would like to thank José Calapez and Dr. Sébastien Chomette from LITEN lab in CEA Grenoble for the training on compressive tests, Nicolas Lemaitre for his help in scaffold cutting, Clément Darche for his help on the Python code, Dr. Pierre-Henri Jouneau from IRIG (CEA Grenoble) for the training on SEM and assistance in imaging, Dr. Matthieu Koepf from DIESE lab in CEA Grenoble for the training on ATR-FTIR, and Dr. Stéphane Lequien from MEM lab in CEA Grenoble for his experiments of SAXS. Dr. Rodolphe Lartizien and Matthieu Olivetto are acknowledged for their expertise in X-ray scoring.

## Notes

### Competing Interest Statement

The authors have declared no competing interest.

