## supporting information file for "3D-printed polymeric scaffolds with optimized architecture to repair a sheep metatarsal critical-size bone defect"

### Garot et al, Supplementary information

#### 1. Supporting Materials & Methods

##### *BMP-2 loading homogeneity*

3D scaffolds of each geometry selected for implantation in a sheep metatarsal critical-size bone defect (Cubic S, Gyroid S, Gyroid L, and Cubic-Gyroid) were 3D-printed by Fused Deposition Modeling (FDM) and film-coated using a dip coating robot. BMP-2 was loaded in the scaffolds using a loading solution containing 5% of BMP-$2^{Rhod}$. Then, scaffolds were transversally cut into 4 to 5 slices, which were deposited in 24-well microplates with 0.15 M NaCl solution. Slices were imaged using an IN Cell Analyzer 2500HS (General Electric, Buc, France) equipped with a 4X objective.

##### *In vitro degradation of scaffolds*

8 scaffolds of each geometry selected for implantation in a sheep metatarsal critical-size bone defect (Cubic S, Gyroid S, Gyroid L, and Cubic-Gyroid) were 3D-printed by FDM, immerged into 40 mL phosphate buffered saline solution (PBS) at pH 7.4, and incubated at 37°C and 5% $CO_2$. 2 scaffolds of each geometry were coated with the polyelectrolyte film using the dipping robot and crosslinked at EDC30. 3 scaffolds without film and 2 film-coated scaffolds of each geometry were used for pH measurements along time to study the eventual release of acidic products. 3 scaffolds of each geometry were used for weight measurement along time. pH measurements were performed on days 0, 1, 2, 3, 7, and every week until $12^{th}$ week for non-coated scaffolds and on days 0, 1, 4, 5, 7, and every week until $12^{th}$ week for film-coated scaffolds. During the pH study, there was no PBS refill or change. There was one control: 40 mL PBS without scaffold. pH variation was calculated by comparing the pH of PBS with scaffold and PBS without scaffold at each time point. Weight measurements were performed on weeks 0, 2, 3, 4, 6, 8, 9, 10, 11, and 12. At each time point, scaffolds were removed from PBS, rinsed with ultrapure water, and dried for 24h before weighting. There was a PBS refill at week 8. At the end of the study, uniaxial compressive tests were performed on all scaffolds except the film-coated scaffolds.

##### *Film coating homogeneity*

3D scaffolds of each geometry selected for implantation in a sheep metatarsal critical-size bone defect (Cubic S, Gyroid S, Gyroid L, and Cubic-Gyroid) were 3D-printed by FDM and film-coated using a dip coating robot. The last deposited layer was labeled with FITC ($PLL^{FITC}$). After their imaging using a fluorescence macroscope, scaffolds were transversally cut into 4 to 5 slices, which were deposited in 24-well microplates in 0.15 M NaCl solution. Slices were imaged using an IN Cell Analyzer 2500HS (General Electric, Buc, France) equipped with a 4X objective.

##### *Film thickness measurement*

Glass coverslips of 14 mm in diameter were coated with 24 bilayers of the polyelectrolyte film using the dip coating robot as previously described [1]. After their coating, the coverslips were crosslinked at EDC30. After crosslinking rinsing, the coverslips were quickly rinsed with ultrapure water, air-dried under a hood, and stored at 4°C. Before imaging the coverslips with a SEM, they were coated with Platinum and scratched with a blade. They were then imaged by SEM (Ultra 55, Zeiss, Oberkochen, Germany).

#### 2. Supporting Figures and Tables

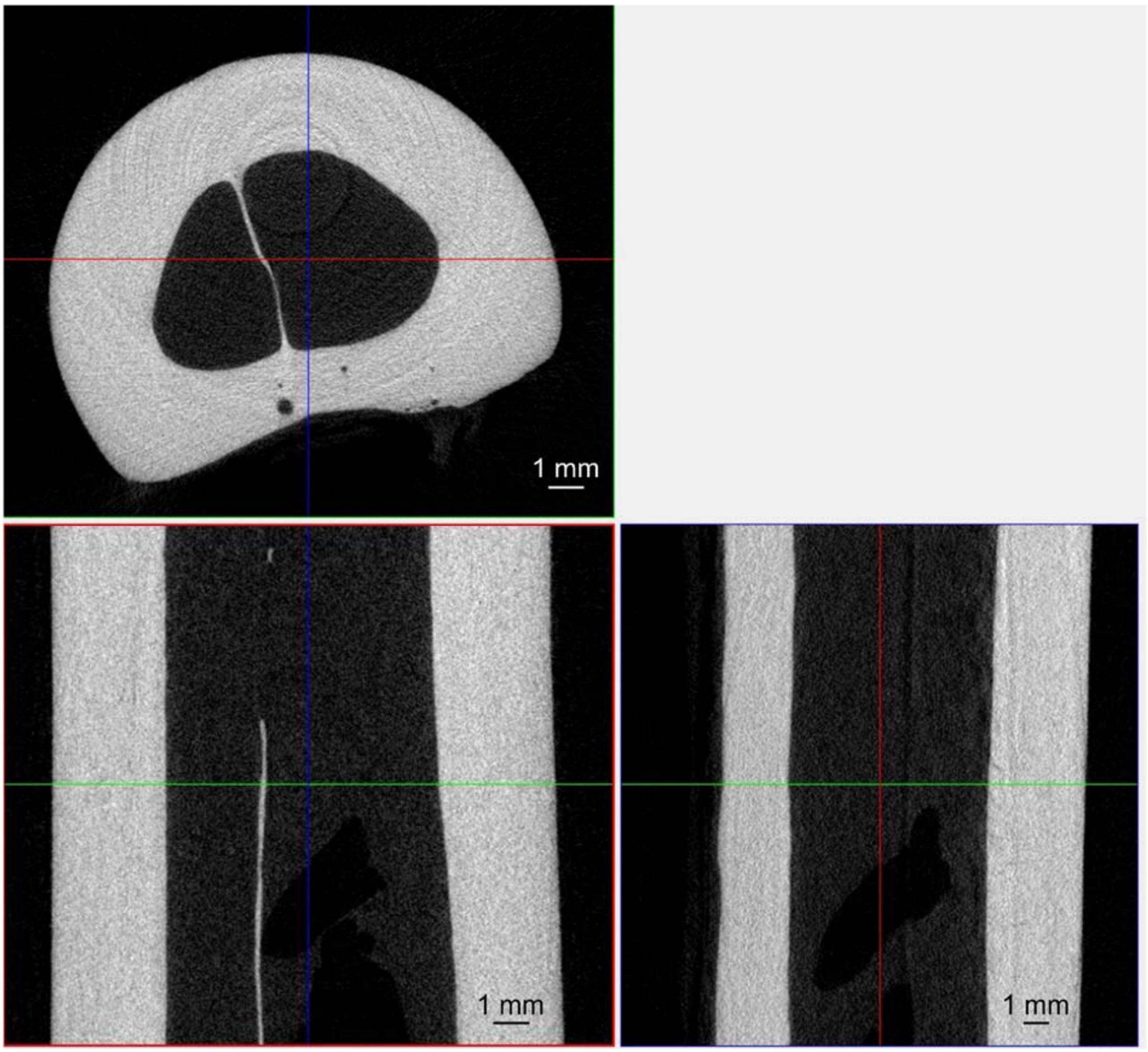


**Figure SI 1. μCT scans of a non-operated sheep metatarsal bone.**

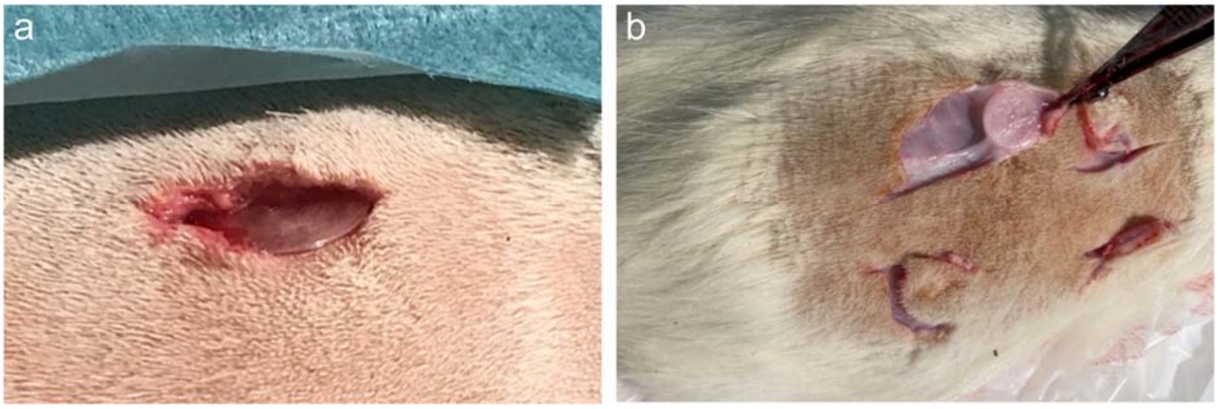


**Figure SI 2. Implantation and explantation of 2D PLA discs in rats.** a) Subcutaneous implantation of 2D PLA discs at day 0. b) Explantation of 2D PLA discs at day 28.

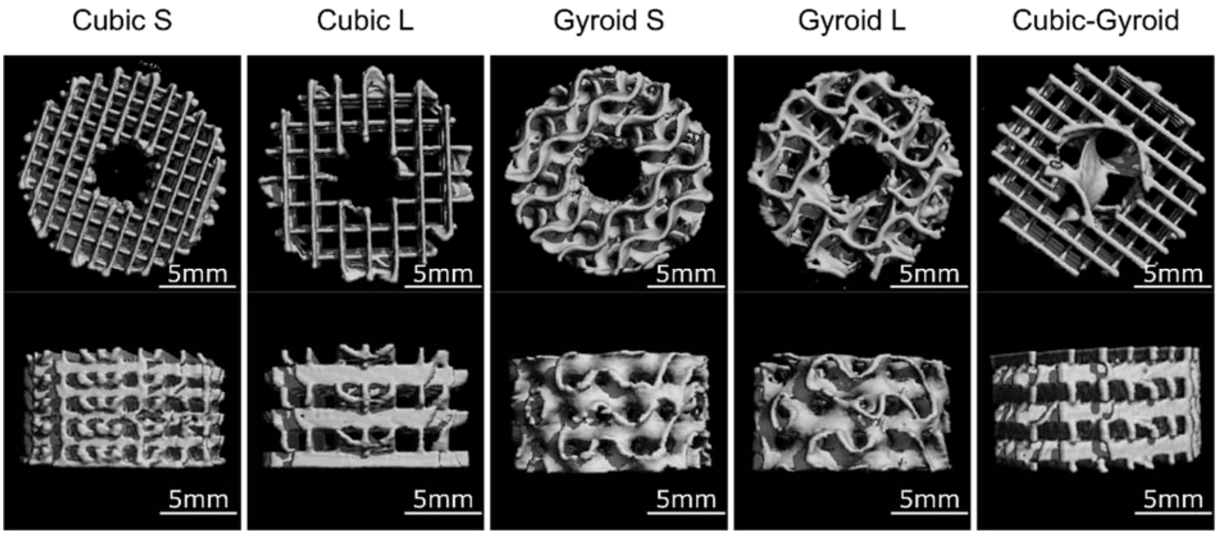


| Unit cell | Surface [cm²] | Porosity [%] | Pore size [μm] |
|---|---|---|---|
| Cubic S | 21 ± 0.5 | 87 ± 1 | 867 ± 31 |
| Cubic L | 14 ± 0.2 | 92 ± 0 | 1337 ± 83 |
| Gyroid S | 16 ± 0.1 | 87 ± 2 | 805 ± 221 |
| Gyroid L | 14 ± 0.4 | 92 ± 1 | 990 ± 263 |
| Cubic-Gyroid | 16 ± 0.2 | 90 ± 0 | 1129 ± 70 |

**Figure SI 3. μCT scans of 3D-printed mini-scaffolds.** Their effective surface, porosity, and pore size are given in the table.

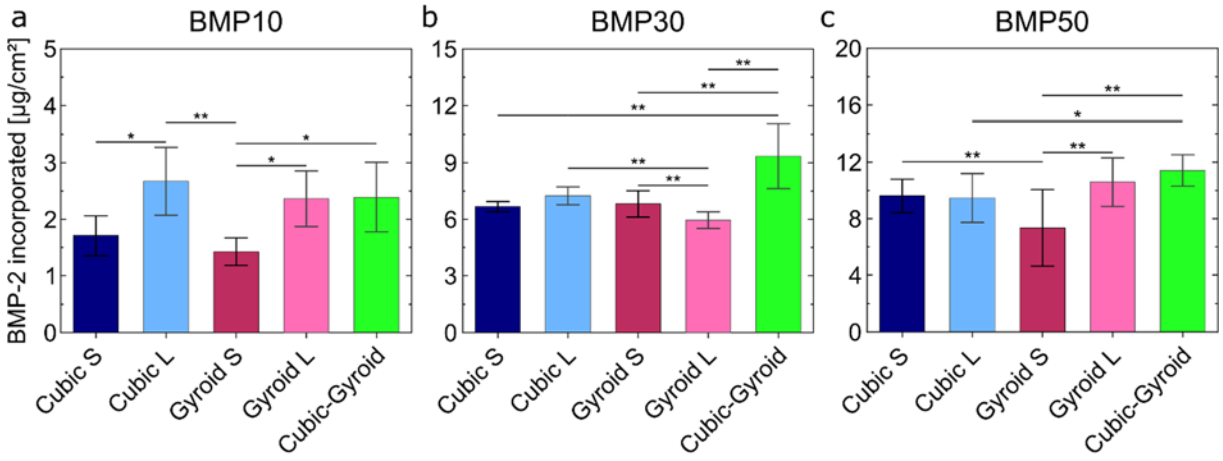


**Figure SI 4. BMP-2 incorporation in 3D mini-scaffolds expressed in surface dose (μg/cm²).** BMP-2 incorporation as a function of scaffold geometry with a loading solution at: a) 10 μg/mL. b) 30 μg/mL. c) 50 μg/mL. *p<0.05; **p<0.01.

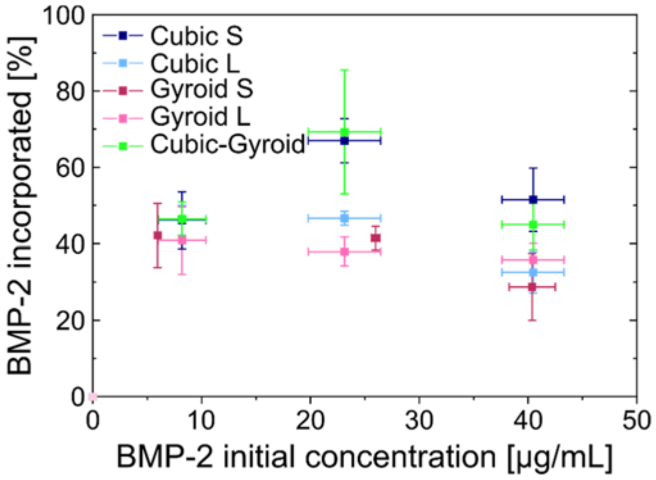


**Figure SI 5. Percentage of BMP-2 incorporated in 3D mini-scaffolds in function of the initial BMP-2 concentration in solution.** This % is given for the studied scaffold geometries: Cubic S and L, Gyroid S and L, Cubic-Gyroid.

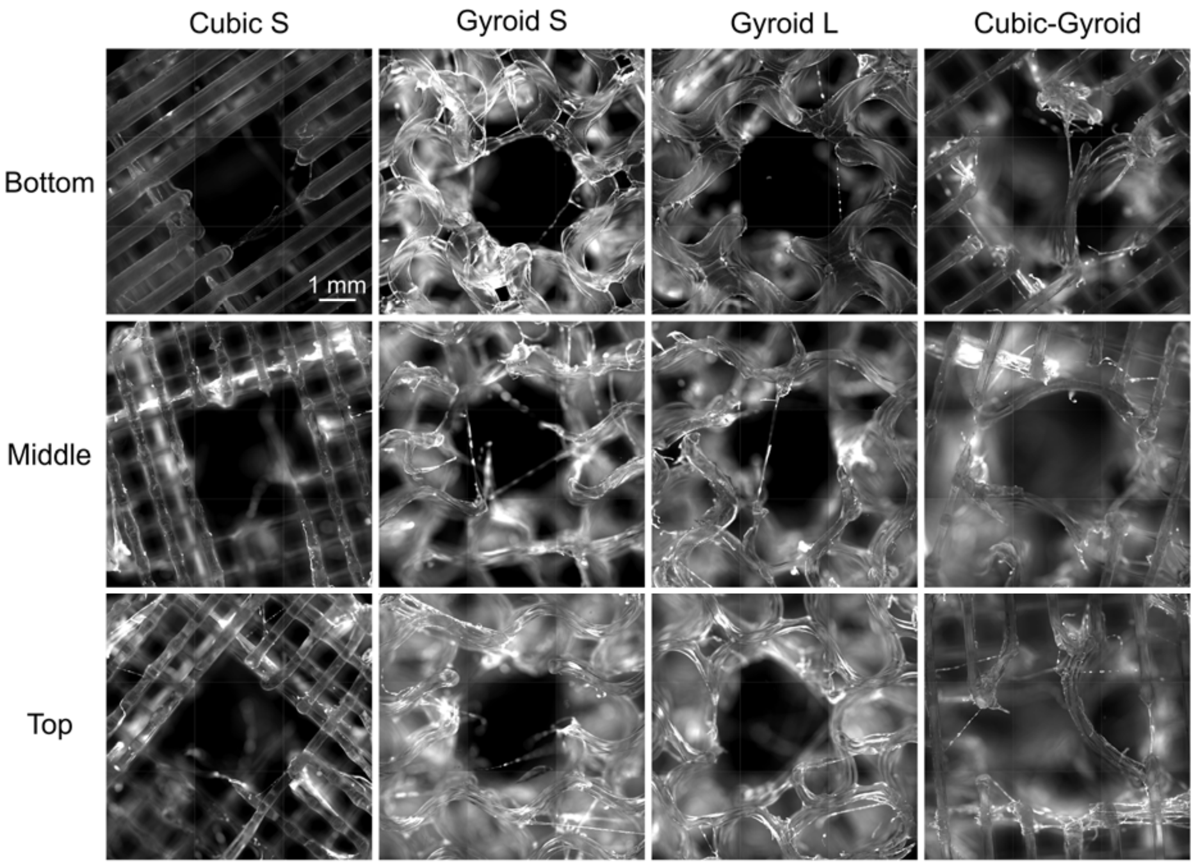


**Figure SI 6. Fluorescence microscopy of 3D scaffolds loaded with BMP-2$^{Rhod}$.** Scale bar is 1 mm for all images.

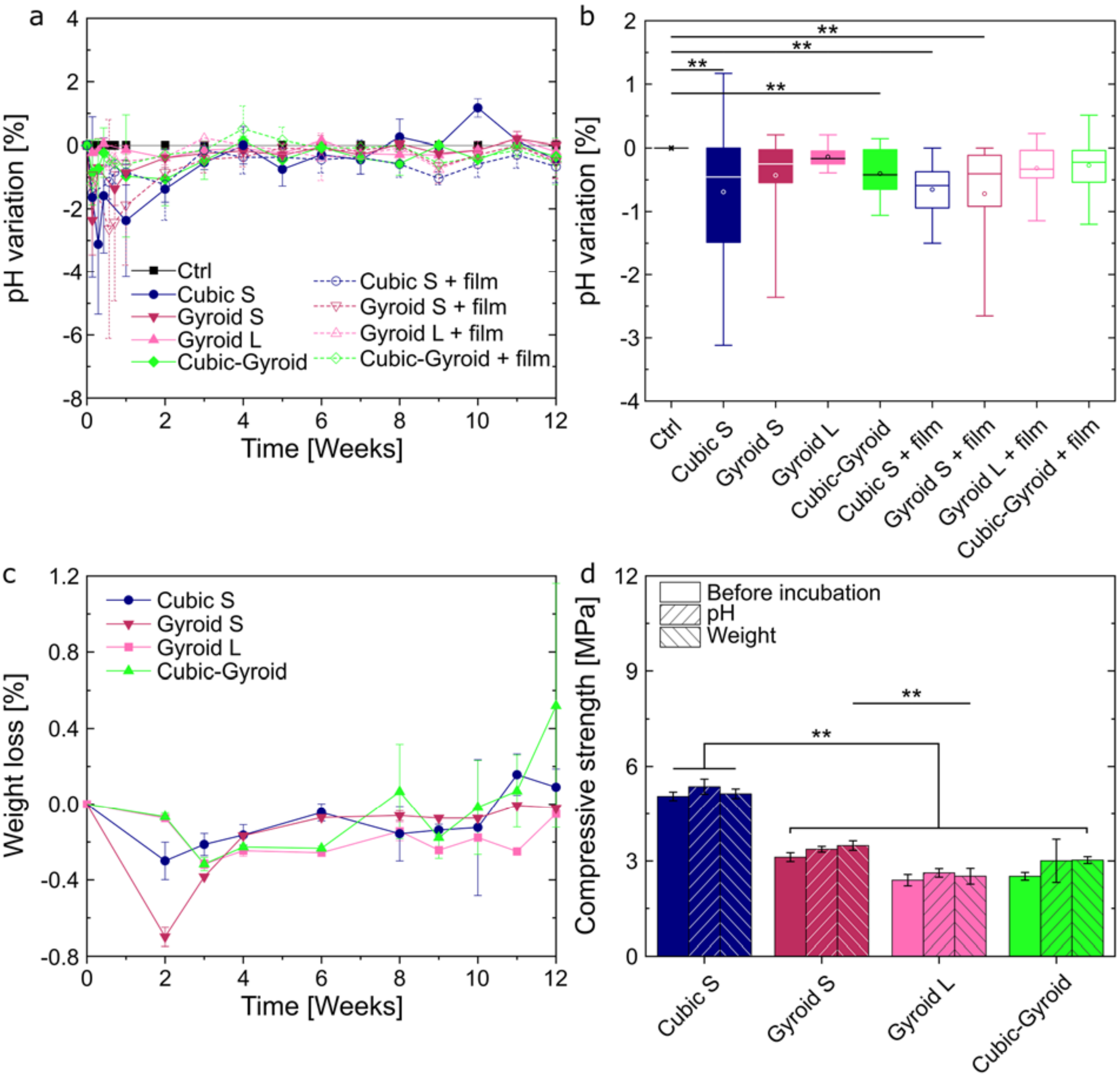


**Figure SI 7. Degradation of 3D PLA scaffolds studied in vitro.** a) pH variation of PBS solution induced by the scaffolds compared to the control (PBS solution without scaffold) as a function of time. b) pH variation of PBS solution induced by the scaffolds compared to the control (PBS solution without scaffold) as a function of the geometry. c) Weight loss of scaffolds as a function of time. d) Compressive strength of scaffolds as a function of the geometry before incubation, after the pH experiment and after the weight experiment. Results are expressed as mean ± SD. *p<0.05 ; **p<0.01.

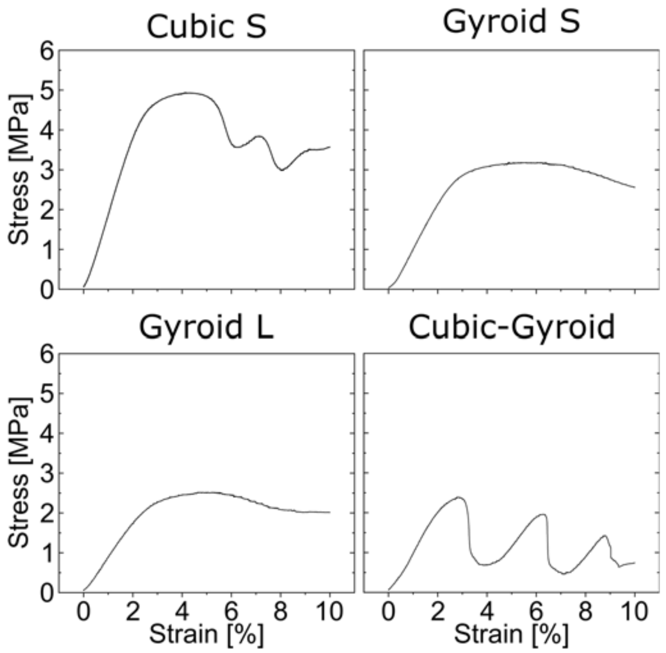


**Figure SI 8. Typical curves representative of uniaxial compressive tests on 3D PLA scaffolds.** Each curve represents the measured stress (MPa) as a function of the strain (expressed in %).

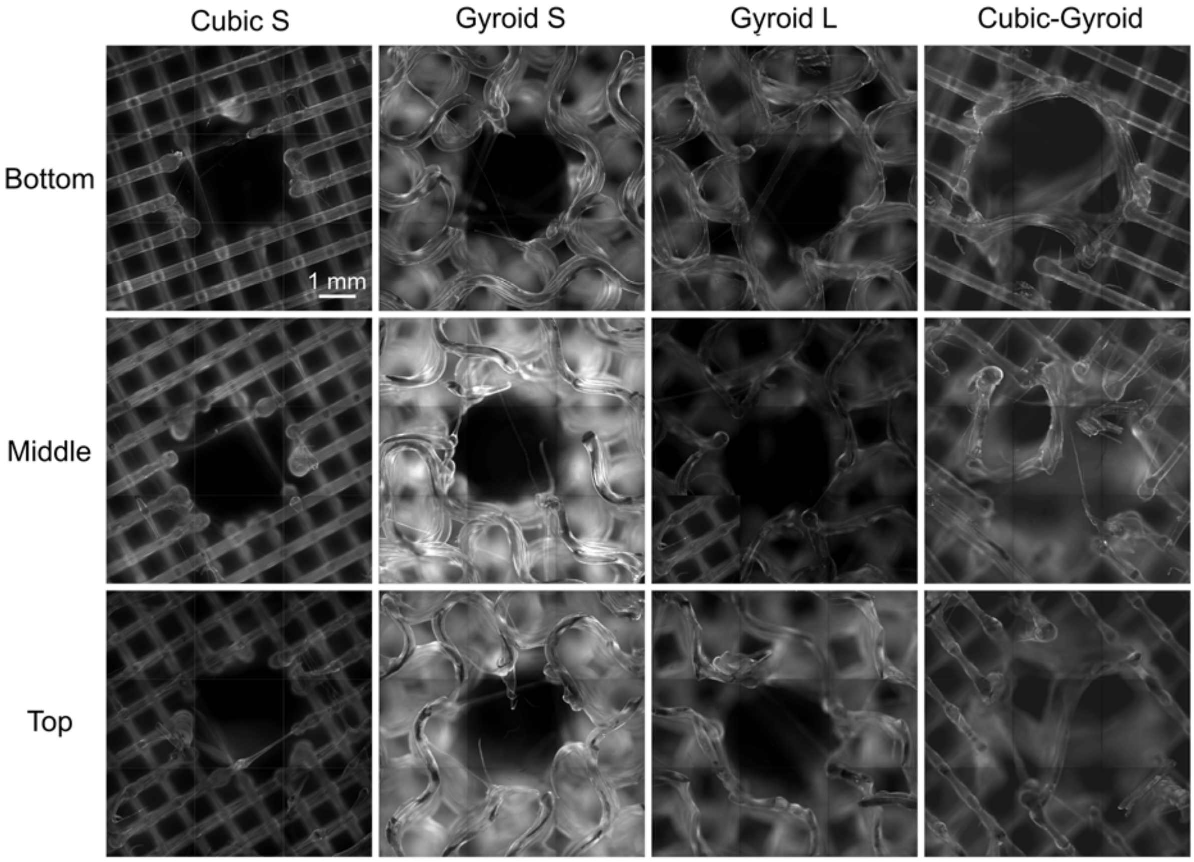


**Figure SI 9. Fluorescence microscopy of 3D scaffolds coated with PLL$^{FITC}$.**
The images are taken at three different positions within the sample: top of the scaffold, middle and bottom. Scale bar is 1 mm for all images.

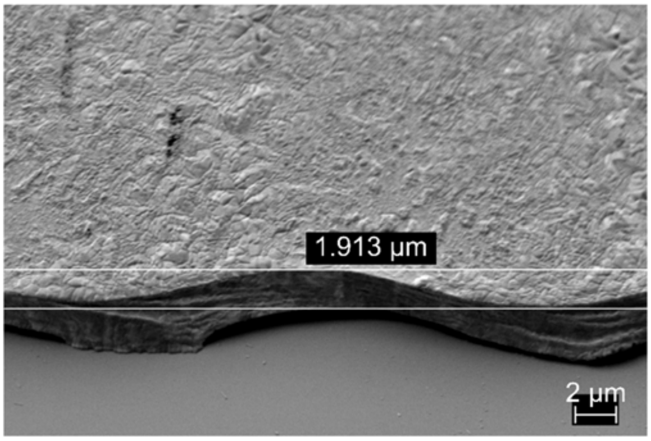


**Figure SI 10. Imaging of a dry film using scanning electron microscopy.**
Imaging of the film was done to measure the film thickness in dry state. It was found to be about 1.9 µm. Scale bare is 2 µm.

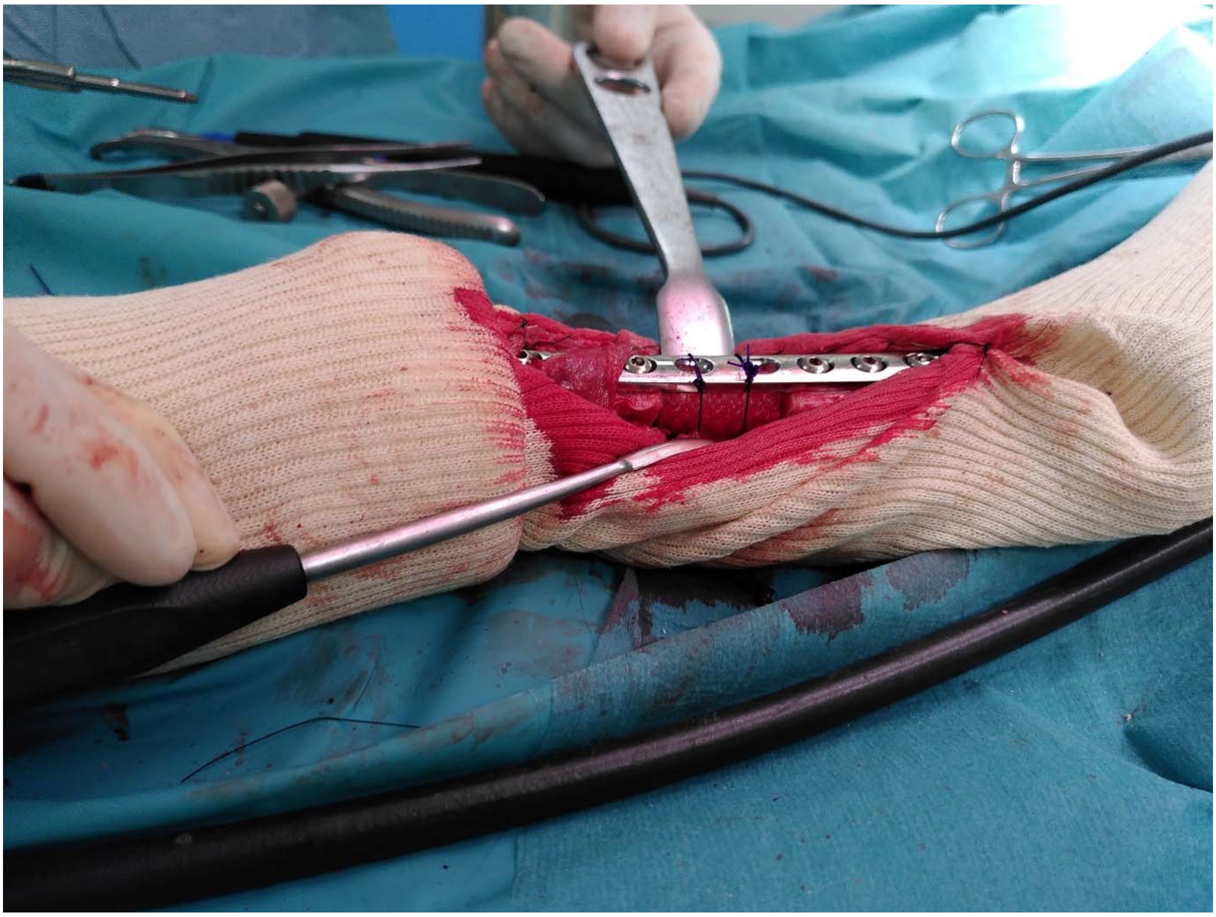

**Figure SI 11. Picture of the implantation of a 3D PLA scaffold in a sheep metatarsal critical-size bone defect.**
First, a critical-size bone defect was created into the sheep metatarsal bone. It was stabilized with an osteosynthesis plate and cortical screws. The scaffold was inserted into the defect and cerclages were used to stabilize the scaffold in contact with the plate.

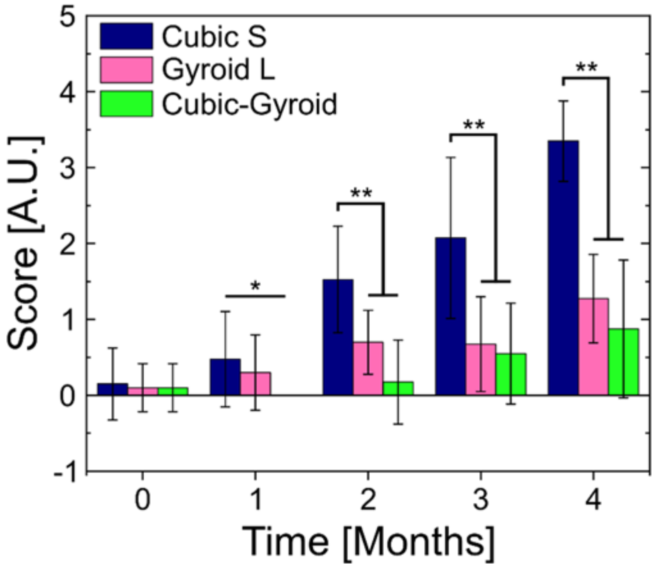


**Figure SI 12. X-ray score as a function of time for the preliminary experiment**. Results are expressed as mean ± SD for all the scores given by 5 clinicians. $*p<0.05$ ; $**p<0.01$.

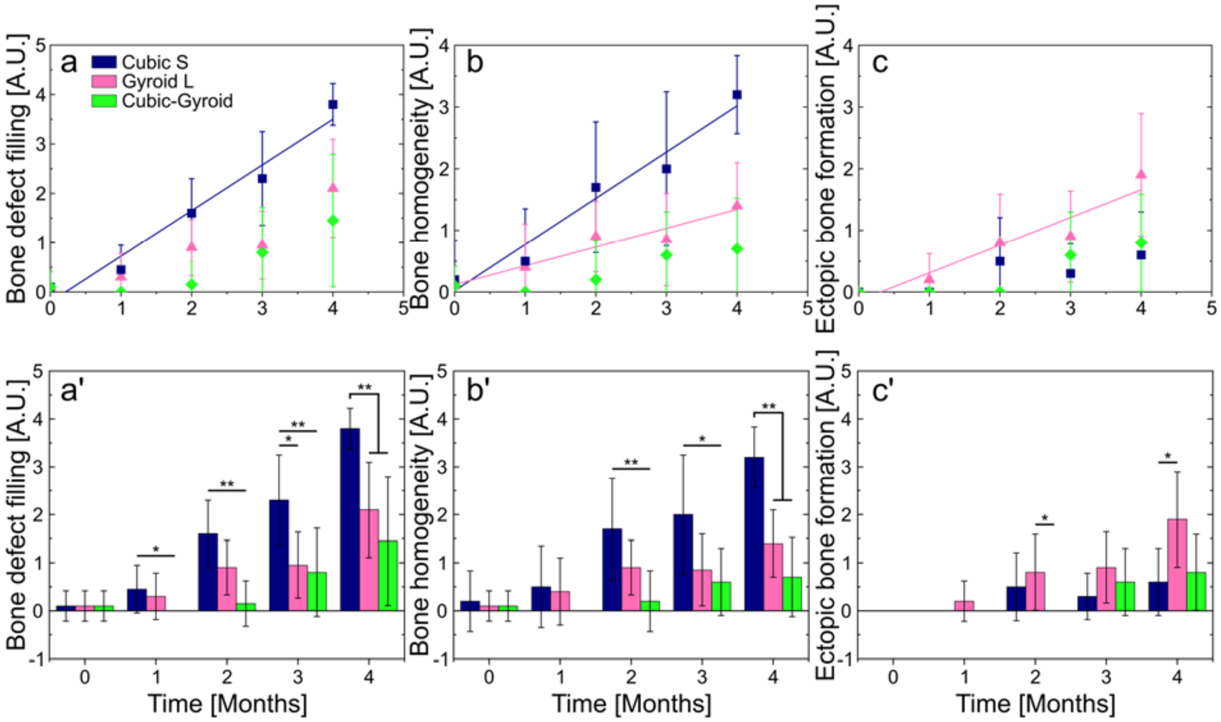


**Figure SI 13. X-ray scores of the preliminary experiment.** The X-ray score were given by five clinicians for each implanted sample a) and a') Bone defect filling as a function of time. b) and b') Bone homogeneity as a function of time. c) and c') Ectopic bone formation as a function of time. Results are expressed as mean ± SD. *$p<0.05$ ; **$p<0.01$.

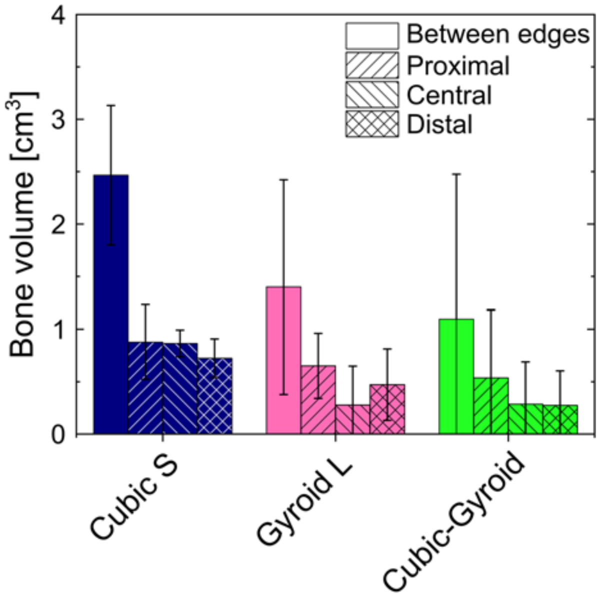


**Figure SI 14. Quantification of bone homogeneity for the preliminary experiment, based on µCT images.**

The bone volume (expressed in $cm^3$) was quantified at different areas along the implant: distal, central, proximal, and between edges/ Results are expressed as mean ± SD for all the studied implants.

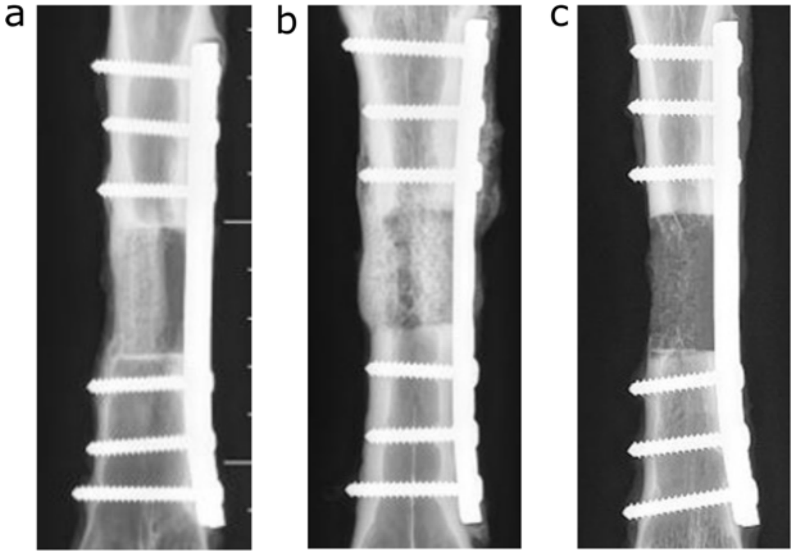


**Figure SI 15. Representative X-ray scans obtained for the different scaffold geometries, whose surface was coated with the biomimetic film loaded with BMP-2.**
a) Cubic S + BMP-2 scaffold that led to partial bridging of the bone defect. b) Gyroid S + BMP-2 scaffold that led to partial bone bridging. c) Gyroid S + BMP-2 scaffold that led to no bone bridging.

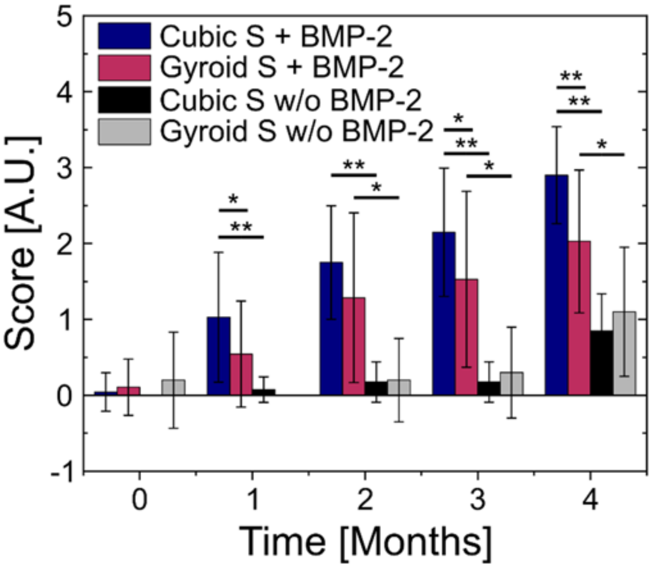


**Figure SI 16. X-ray score as a function of time for the main experiment.** The scores were obtained for the two different scaffold geometries (Cubic C and Gyroid S) in the absence of BMP-2 (w/o BMP-2) in the biomimetic film or in the presence of BMP-2 in the biomimetic film (+ BMP-2). The results are expressed as mean ± SD for all sample studied, and for each sample, independent scoring by 5 clinicians (see . *p<0.05 ; **p<0.01.

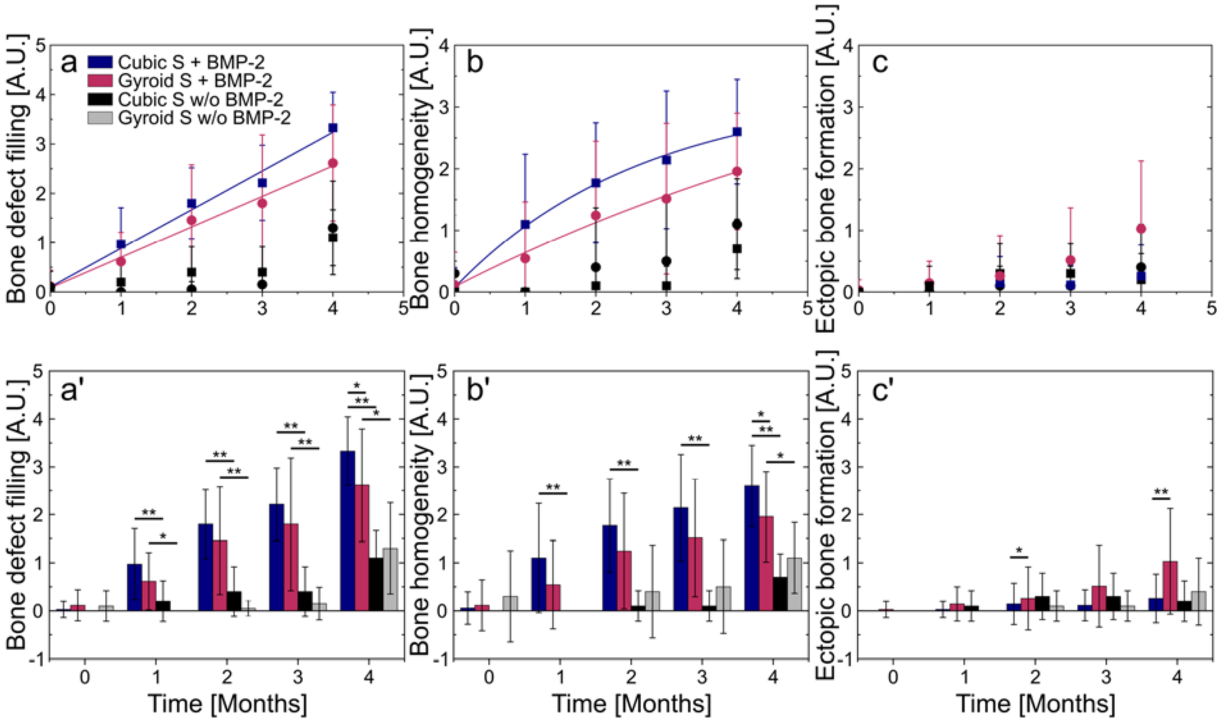


**Figure SI 17. Parameters deduced from X-ray scores for the Cubic S and Gyroid S geometries studied during the main experiment.** Bone defect filling, bone homogeneity and ectopic bone formation were quantified by the clinicians for each sample, and for each time point**.** Data are represented either as mean value $\pm$ SD as data point (upper row) or as bar plots (bottom row, prime plots) as a function of time, up to four months. a) and a') Bone defect filling. b) and b') Bone homogeneity. c) and c') Ectopic bone formation. The fit of the data are also given. *$p<0.05$ ; **$p<0.01$.

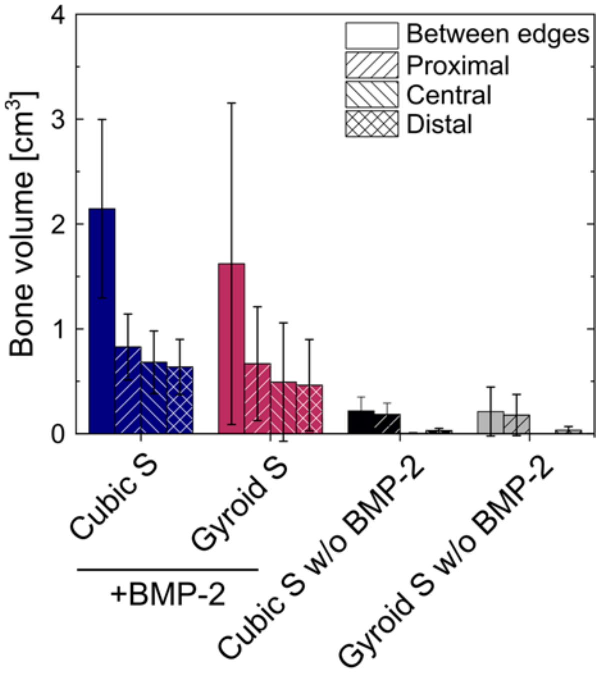


**Figure SI 18. Bone homogeneity quantification for the main experiment, based on µCT images.** The bone volume (expressed in $cm^3$) was quantified at different areas along the implant: distal, central, proximal, and between edges/ Results are expressed as mean ± SD for all the studied implants.

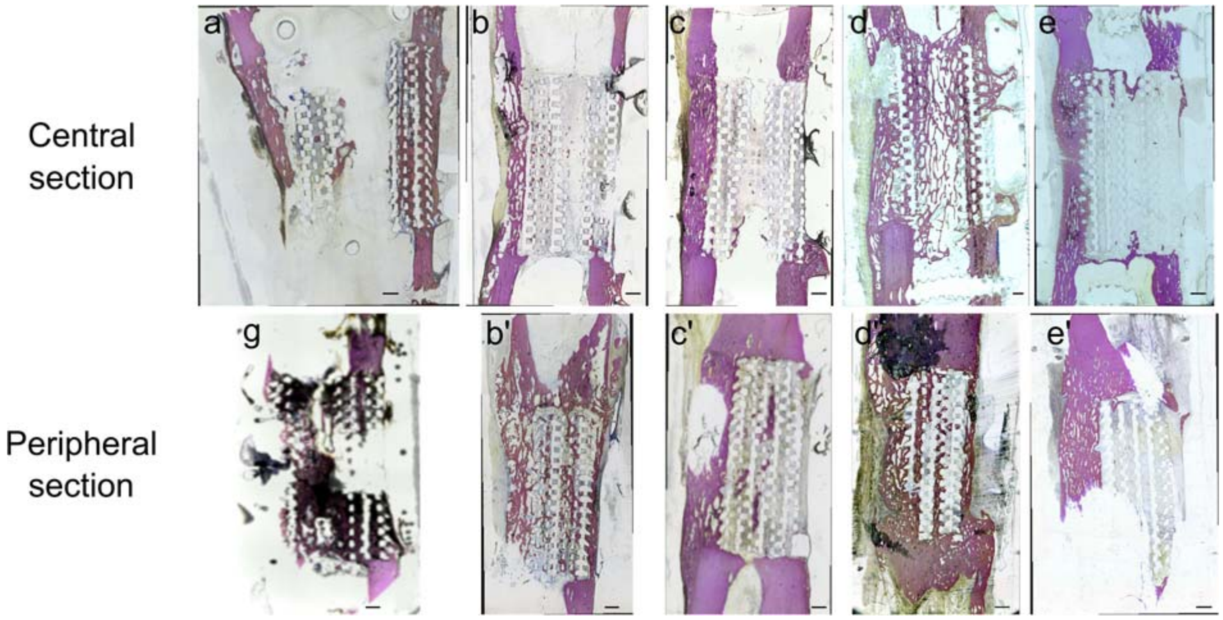


**Figure SI 19. Histological sections of the whole explants of the Cubic S + BMP-2 group.** a) and g) represent the two Cubic S scaffolds from the preliminary experiment. b)-e') represent the Cubic S + BMP-2 scaffolds from the main experiment. Scale bar is 2 mm.

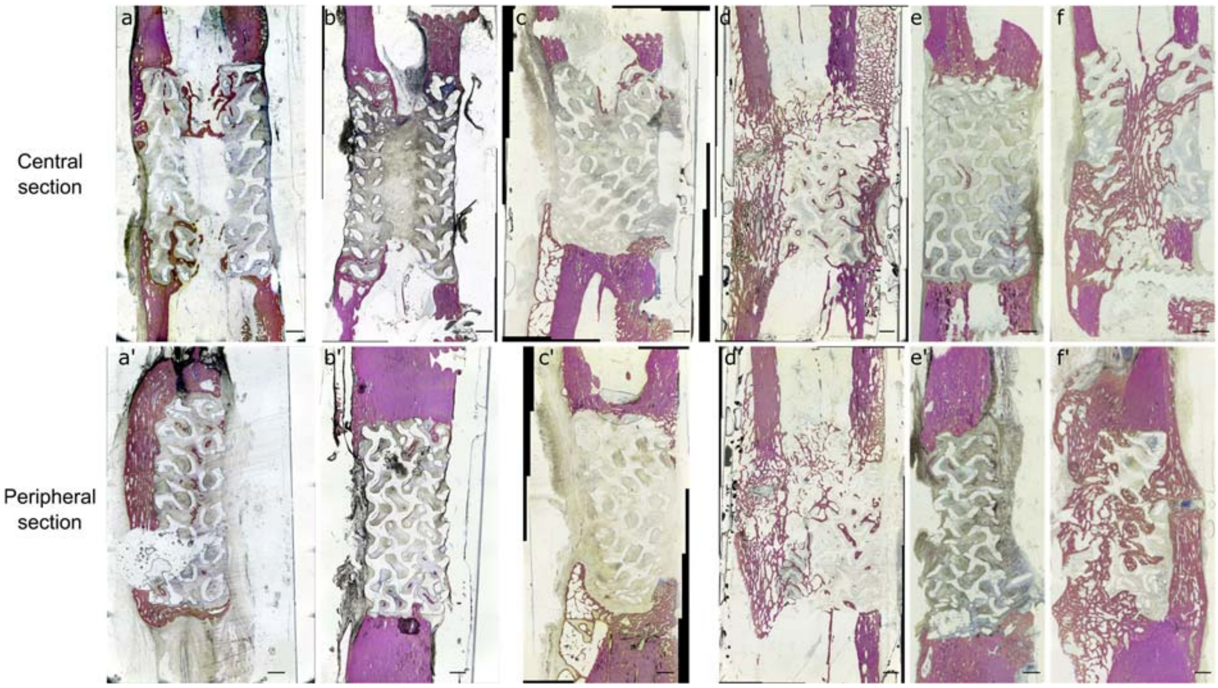


**Figure SI 20. Histological sections of a whole explant of the Gyroid S + BMP-2 group.** Scale bar is 2 mm.

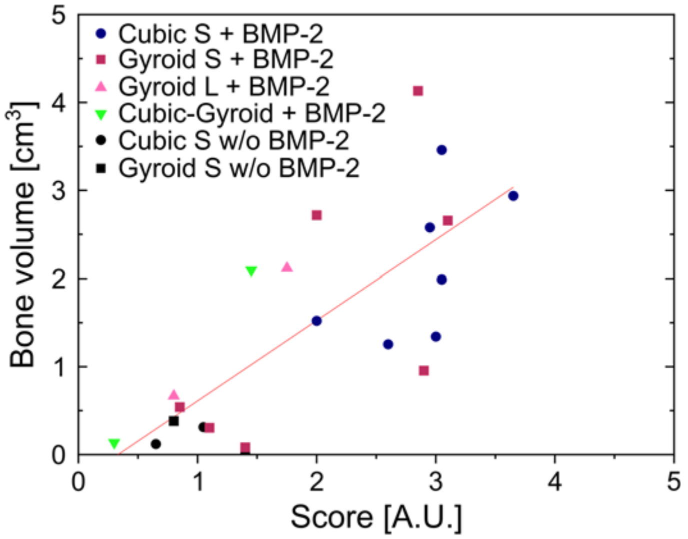


**Figure SI 21. Correlation between the qualitative X-ray score given by clinicians and the quantitative measurement of the bone volumes using computed tomography.** The bone volume (cm3) of each sample is represented as a funciton of the mean score given by the clinicians based on X-ray imaging (at 4 months). Each type of scaffold is represented by a given symbol and color. Data corresponding to all samples are pooled on the same graph. Results are expressed as mean ± SD. The line is linear fit of the data.

**Table SI 1. 3D-printed scaffolds for the design of experiment.** O: outer ring; I: inner ring.

| Center of the cylinder | | Pore shape | Infill density (%) | Number of scaffolds |
|---|---|---|---|---|
| Pattern | Diameter (mm) | | | |
| Hole | 5 | Cubic | 33 | 3 |
| Hole | 5 | Cubic | 41.5 | 3 |
| Hole | 5 | Cubic | 50 | 3 |
| Hole | 5 | Gyroid | 15 | 3 |
| Hole | 5 | Gyroid | 25 | 3 |
| Hole | 5 | Gyroid | 35 | 3 |
| Gyroid | 5 | O: Cubic<br>I: Gyroid | O: 44.4<br>I: 35 | 1 |
| Gyroid | 5 | O: Cubic<br>I: Gyroid | O: 44.4<br>I: 10 | 1 |
| Gyroid | 5 | O: Cubic<br>I: Gyroid | O: 16.7<br>I: 35 | 1 |
| Gyroid | 5 | O: Cubic<br>I: Gyroid | O: 16.7<br>I: 10 | 1 |
| Gyroid | 5 | O: Zigzag<br>I: Gyroid | O: 44.4<br>I: 35 | 1 |
| Gyroid | 5 | O: Zigzag<br>I: Gyroid | O: 44.4<br>I: 10 | 1 |
| Gyroid | 5 | O: Zigzag<br>I: Gyroid | O: 16.7<br>I: 35 | 1 |
| Gyroid | 5 | O: Zigzag<br>I: Gyroid | O: 16.7<br>I: 10 | 1 |
| Gyroid | 7 | O: Cubic opened<br>I: Gyroid | O: 44.4<br>I: 35 | 1 |
| Gyroid | 7 | O: Cubic opened<br>I: Gyroid | O: 44.4<br>I: 10 | 1 |
| Gyroid | 7 | O: Cubic opened<br>I: Gyroid | O: 16.7<br>I: 35 | 1 |
| Gyroid | 7 | O: Cubic opened<br>I: Gyroid | O: 16.7<br>I: 10 | 1 |
| Gyroid | 7 | O: Zigzag<br>I: Gyroid | O: 44.4<br>I: 35 | 1 |
| Gyroid | 7 | O: Zigzag<br>I: Gyroid | O: 44.4<br>I: 10 | 1 |
| Gyroid | 7 | O: Zigzag<br>I: Gyroid | O: 16.7<br>I: 35 | 1 |
| Gyroid | 7 | O: Zigzag<br>I: Gyroid | O: 16.7<br>I: 10 | 1 |
| Gyroid | 7 | O: Cubic opened<br>I: Gyroid | O: 24.2<br>I: 22.5 | 3 |
| Gyroid | 7 | O: Zigzag | O: 24.2 | 3 |

**Table SI 2. Fitting characteristics of the curves from Figure 4a and b.** a) Fitting characteristics of the curves from **Figure 4a**. b) Fitting characteristics of the curves from **Figure 4b**. For both tables, the fit of Cubic S, Cubic L, Gyroid S, and Cubic-Gyroid was an exponential fit following the equation $y = y_0 + A_1 * e^{-x/k}$. For Gyroid L, the fit was linear following the equation $y = a + b * x$.

a)

| Geometry | Fitting parameters | | | $R^2$ | Adjusted $R^2$ |
|---|---|---|---|---|---|
| | $y_0$[μg] | $A_1$[μg] | k[μg/mL] | | |
| Cubic S | 450.4 ± 408.7 | -456.3 ± 401.4 | 63.7 ± 77.7 | 0.99 | 0.97 |
| Cubic L | 178.9 ± 37.1 | -180.9 ± 35.2 | 29.7 ± 11.3 | 1 | 0.986 |
| Gyroid S | 146.5 ± 48.8 | -151.1 ± 45.4 | 21.8 ± 16.4 | 0.98 | 0.945 |
| Cubic-Gyroid | 256.1 ± 141.3 | -263.2 ± 134.1 | 30.9 ± 30.2 | 0.97 | 0.919 |
| Gyroid L | a[μg]=1.1 ± 1.6<br>b[mL]=3.5 ± 0.1 | | | 1 | 0.999 |

b)

| Geometry | Fitting parameters | | | $R^2$ | Adjusted $R^2$ |
|---|---|---|---|---|---|
| | $y_0$[μg/cm²] | $A_1$[μg/cm²] | k[μg/mL] | | |
| Cubic S | 21.1 ± 19.1 | -21.3 ± 18.8 | 63.7 ± 77.7 | 0.99 | 0.97 |
| Cubic L | 12.9 ± 2.7 | -13.1 ± 2.5 | 29.7 ± 11.3 | 1 | 0.99 |
| Gyroid S | 9.1 ± 3.0 | -9.4 ± 2.8 | 21.8 ± 16.4 | 0.98 | 0.95 |
| Cubic-Gyroid | 16.3 ± 9.0 | -16.8 ± 8.5 | 30.9 ± 30.2 | 0.97 | 0.92 |
| Gyroid L | a[μg]=1.11 ± 1.57<br>b[mL]=3.54 ± 0.07 | | | 1 | 1 |

**Table SI 3. Identification of the peaks obtained using infrared spectroscopy (ATR-FTIR mode).**

| Peak number | IR frequencies ($cm^{-1}$) | Assignments | Remarks |
|---|---|---|---|
| 1 | 3100-3700 | O-H stretching absorbance [2,3] | |
| 2 | 2997 | $\nu_{as}CH_3$ (-CH- stretch) [4–9] | |
| | 2946 | $\nu_{s}CH_3$ (-CH- stretch) [4–9] | Peak only present in normal PLA spectra |
| | 2875 | νCH [4] | Peak only present in normal PLA spectra |
| 3 | 1755 | ν(C=O) (-C=O stretching vibration) [4–10] | |
| 4 | 1650 | | Bending vibration from water [3,9] |
| 5 | 1455 | $\delta_{as}CH_3$ (-$CH_3$ bend) [3,4,7,8] | |
| | 1384 | $\delta_{s}CH_3$ (-CH- bend) [4,7,8] | Characteristic peak of amorphous PLA |
| | 1368 | $\delta_1CH + \delta_sCH_3$ [4,8] | Peak only present in medical-grade PLA spectra and characteristic of semicrystalline PLA |
| | 1363 | $\delta_1CH + \delta_sCH_3$ (-CH- bend) [4,7] | Peak only present in normal PLA spectra and characteristic of amorphous PLA |
| | 1360 | $\delta_1CH + \delta_sCH_3$ [4] | Peak only present in medical-grade PLA spectra and characteristic of semicrystalline PLA |
| | 1305 | $\delta_2CH$ [4,8] | |
| 6 | 1267 | δCH + νCOC<br>νCH+C(C-CO-O) (-C=O bend) [4,7,8,10] | Weaker peak for medical-grade PLA. The decrease of absorptivity is characteristic of PLA crystallization |
| | 1213 | $\nu_{as}COC + r_{as}CH_3$ [4,10] | Peak only present in medical-grade PLA spectra and characteristic of semicrystalline PLA |
| | 1210 | $\nu_{as}COC + r_{as}CH_3$ [4] | Peak only present in normal PLA spectra and characteristic of amorphous PLA. Weak band |

| | | | |
|---|---|---|---|
| | 1184 | $\nu_{as}COC + r_{as}CH_3$ (-C-O- stretch) [4,6–10] | |
| | 1130 | $r_{as}CH_3$<br>$r_sCH_3$<br>(-C-O- stretch) [4,7,8,10] | |
| | 1089 | $\nu_sCOC$<br>$\nu_{as}$(O-C-CO)<br>(-C-O- stretch) [4,7,8,10] | |
| 7 | 1046 | $\nu(C\text{-}CH_3)$ [4,8,10]<br>-OH bend [7] | |
| 8 | 956 | $rCH_3 + \nu CC$ [4,6,8,10] | Very weak peak for medical-grade PLA. The intensity of the band decreases with crystallization |
| | 925 | $rCH_3 + \nu CC$ (-C-C stretch) [4,7,10] | Peak only present in medical-grade PLA spectra and characteristic of semicrystalline PLA (characteristic of α crystals |
| 9 | 871 | νC-COO (-C-C- amorphous phase) [4,6–8] | Characteristic peak of amorphous PLA |
| 10 | 755 | δC=O (-C-C crystalline phase) [4,6,8] | Very weak peak |
| | 740 | δC=O [4] | |
| | 708 [8] | | |

**Table SI 4. Identification of the diffraction peaks obtained by X-ray analysis of the PLA samples.** Here, cobalt radiation (λ=0.17903 nm) was used. Acquisitions are made in θ/2θ reflexion mode. hkl are the Miller indices.

| | | | Normal PLA | | Medical-grade PLA | |
|---|---|---|---|---|---|---|
| h | k | l | 2θ [°] | Interplanar distance [Å] | 2θ [°] | Interplanar distance [Å] |
| 2 | 0 | 0 | 17.96 | 5.74 | 18.98 | 5.43 |
| 2 | 0 | 3 | - | - | 21.66 | 4.77 |
| 2 | 1 | 1 | 24.88 | 4.16 | 25.82 | 4.01 |
| 0 | 0 | 8 | - | - | 28.30 | 3.66 |
| 3 | 1 | 0 | - | - | 33.28 | 3.13 |
| 0 | 2 | 5 | 37.62 | 2.78 | 38.02 | 2.75 |
